# Gut metabolite L-lactate supports *Campylobacter jejuni* population expansion during acute infection

**DOI:** 10.1101/2023.10.02.560557

**Authors:** Ritam Sinha, Rhiannon M. LeVeque, Sean M. Callahan, Shramana Chatterjee, Nejc Stopnisek, Matti Kuipel, Jeremiah G. Johnson, Victor J. DiRita

## Abstract

How the microaerobic pathogen *Campylobacter jejuni* establishes its niche and expands in the gut lumen during infection is poorly understood. Using six-week-old ferrets as a natural disease model, we examined this aspect of *C. jejuni* pathogenicity. Unlike mice, which require significant genetic or physiological manipulation to become colonized with *C. jejuni*, ferrets are readily infected without the need to disarm the immune system or alter the gut microbiota. Disease after *C. jejuni* infection in ferrets reflects closely how human *C. jejuni* infection proceeds. Rapid growth of *C. jejuni* and associated intestinal inflammation was observed within two-three days of infection. We observed pathophysiological changes that were noted by cryptic hyperplasia through the induction of tissue repair systems, accumulation of undifferentiated amplifying cells on the colon surface, and instability of HIF-1α in colonocytes, which indicated increased epithelial oxygenation. Metabolomic analysis demonstrated that lactate levels in colon content were elevated in infected animals. A *C. jejuni* mutant lacking *lctP*, which encodes an L-lactate transporter, was significantly decreased for colonization during infection. Lactate also influences adhesion and invasion by *C. jejuni* to a colon carcinoma cell line (HCT116). The oxygenation required for expression of lactate transporter (*lctP*) led to discovery of a putative thiol based redox switch regulator (LctR) that may repress *lctP* transcription under anaerobic conditions. Our work provides new insights into the pathogenicity of *C. jejuni*.

**Significance:** There is a gap in knowledge about the mechanisms by which *C. jejuni* populations expand during infection. Using an animal model which accurately reflects human infection without the need to alter the host microbiome or the immune system prior to infection, we explored pathophysiological alterations of the gut after *C. jejuni* infection. Our study identified the gut metabolite L-lactate as playing an important role as a growth substrate for *C. jejuni* during acute infection. We identified a DNA binding protein, LctR, that binds to the *lctP* promoter and may repress *lctP* expression, resulting in decreased lactate transport under low oxygen levels. This work provides new insights about *C. jejuni* pathogenicity.

## Introduction

Foodborne diseases are important threats in both developed and developing countries. Among the more prevalent foodborne bacterial pathogens is *Campylobacter jejuni*, annually responsible for an estimated 1.5 million cases of gastroenteritis (1). With the emergence of antibiotic resistant strains, the choice of antibiotic to treat these infections may soon be limited; the U.S. Centers for Disease Control and Prevention considers drug resistant *Campylobacter* infection as a serious threat (2, 3). Common symptoms of gastroenteritis mediated by *C. jejuni* are diarrhea (sometimes bloody), fever, vomiting and abdominal cramps. Symptoms typically begin within two to five days after infection and last about a week. *C. jejuni* gastroenteritis can be a more persistent, and even life-threating, infection in immunocompromised patients such as those with AIDS or hypogammaglobulinemia (3, 4). It can also have the serious post-infection sequelae of Guillain-Barré syndrome (5). The precise mechanisms of its pathogenicity remain uncertain as *C. jejuni* does not encode pathogenicity islands associated with secretion of toxins or other effectors that would enable it to manipulate host cell biology or survive intracellularly, as do other gastrointestinal pathogens such as *Salmonella*, *Shigella*, and *Vibrio cholerae* (4, 5). More well-studied pathogens such as facultative anaerobes in the family *Enterobacteriaceae*, including *Citrobacter rodentium* and *Salmonella typhimurium*, induce crypt colonic hyperplasia via virulence factors, and regulate mucosal epithelial oxygenation and lactate concentration in the gut during inflammation, which influence their growth and pathogenicity (6–8).

Lack of an ideal animal model means there remain gaps in understanding the precise mechanism of *C.jejuni* disease progression *in vivo*. Some questions regarding *C.jejuni* pathogenicity that remain unanswered are: i) How does *C.jejuni* establish its niche and expand in the gut lumen during the inflammatory stage of infection? ii) How does *C. jejuni* compete with the gut microbiota for colonization in gut? iii) What is the major carbon source for *C.jejuni* growth during acute infection?

Our main goal is to understand environmental conditions during infection that influence colonization and population expansion of *C.jejuni*. We used a ferret-infection model to investigate pathogenicity of *C. jejuni*. This model has several advantages over currently used mouse models, including no required antibiotic treatment or other manipulation of the gut microbiota prior to infection (9), or disarming key components of the immune system such as MyD88 (10), NF-kB (11), or IL-10 (12). Unlike ferrets, these knockout mouse models do not faithfully mimic the signs or intestinal pathology of human campylobacteriosis.

In this study, we assessed the disease dynamics of *C. jejuni* in five-to-six-week-old ferrets, investigating colonization, localization patterns, and pathophysiological changes in the gut during early and acute stage of infection after orogastric inoculation with *C. jejuni.* We describe changes of gut microbiota and metabolites during the acute (inflammatory) stage of infection. We identified L-lactate as a potential carbon source for growth, and we demonstrate that the lactate transporter operon (*lctP*) is essential for *C. jejuni* expansion and colonization in the inflamed gut. Finally, we identified a potential redox switch regulator (LctR) of *lctP* which, we hypothesize, in its oxidized state comes off the *lctP* promoter enabling lactate uptake and growth in the inflamed gut.

## Results

### *C. jejuni* grows in the ferret gut during inflammation

Experimental infection of humans and primates with *C. jejuni* have used oral doses as high as 10^9^ to 10^10^ CFU to achieve infection and acute illness, although infection may occur with much lower doses during outbreaks (1, 2). To test dose-dependent infection by *C. jejuni* in ferrets, aged five-to-six weeks old animals were infected with two different doses, 10^8^ or 10^9^ CFU/ml, of *C.jejuni* strain 11168 and *C. jejuni* loads in feces were determined and pathophysiological changes in the colon were scored by histological analysis. At the higher dose of infection animals shed approximately 10^7^ CFU/gram of stool after 24 hours, as compared to 5 x 10^4^ CFU/gram of stool from animals infected at the lower dose. Animals infected at the higher dose shed approximately 10^7^ CFU/gram of stool after 24 hours, as compared to 5 x 10^4^ CFU/gram of stool from animals infected at the lower dose. By 72 hours, animals infected at the higher dose exhibited increased shedding of *C. jejuni*, to nearly 10^10^ CFU/gram of stool, while those infected at the lower dose were shedding only slightly higher numbers of *C. jejuni* than they were at 24 hours (no *C. jejuni* were detected in the PBS control group after 72h of post infection; **FigS1A**). Comparative histopathological scoring analysis and H and E staining at 72h post-infection demonstrated no histopathological differences between the PBS and the 10^8^ groups but moderate to severe gastroenteritis signs in the 10^9^ group (epithelial cell damage, inflammatory cell infiltration, goblet cell depletion, cryptic hyperplasia and cryptic abscess) **(Fig S1B,C).** Analysis with anti-*Campylobacter* antibody identified many *C. jejuni* near colonic epithelial cells in animals infected with the higher dose, whereas *C. jejuni* was present only within the colonic lumen area in animals infected with the lower dose **(Fig S1B).** These findings indicate that inoculation with 10^9^ CFU of *C. jejuni* 11168 causes infection and acute gastroenteritis after 72 hours in ferrets, which is like what is observed for human campylobacteriosis (1, 3). These data also suggest that intestinal inflammation supports *C. jejuni* expansion in ferret gut, similar to how intestinal inflammation induces growth of facultative anaerobes such as *E. coli*, *Salmonella enterica sv.* Typhimurium and *Citrobacter rodentium* (4, 5).

We investigated other pathophysiological changes during the acute stage of *C. jejuni* infection to explore potential determinants of its growth. We infected ferrets with 5×10^9^ CFU/ml of *C. jejuni* strain 11168 and measured bacterial levels in colon, small intestine, mesenteric lymph node (MLN), liver, and spleen days one and three post-infection. Levels of *C. jejuni* at every site were higher on day three post-infection compared to day one, with the highest concentration in the colon, where *C. jejuni* levels rose from approximately 10^6^ cfu/gm of tissue on day one to approximately 10^10^ cfu/gm on day three (**Fig 1A**). Expansion of *C. jejuni* in the colon on day three was also confirmed by 16sRNA sequencing (**Fig 1B**). Although we did not explore it further, the presence of *C. jejuni* in deeper tissue such as liver and spleen on day three is evidence of systemic infection in the ferret.

**Figure 1:**
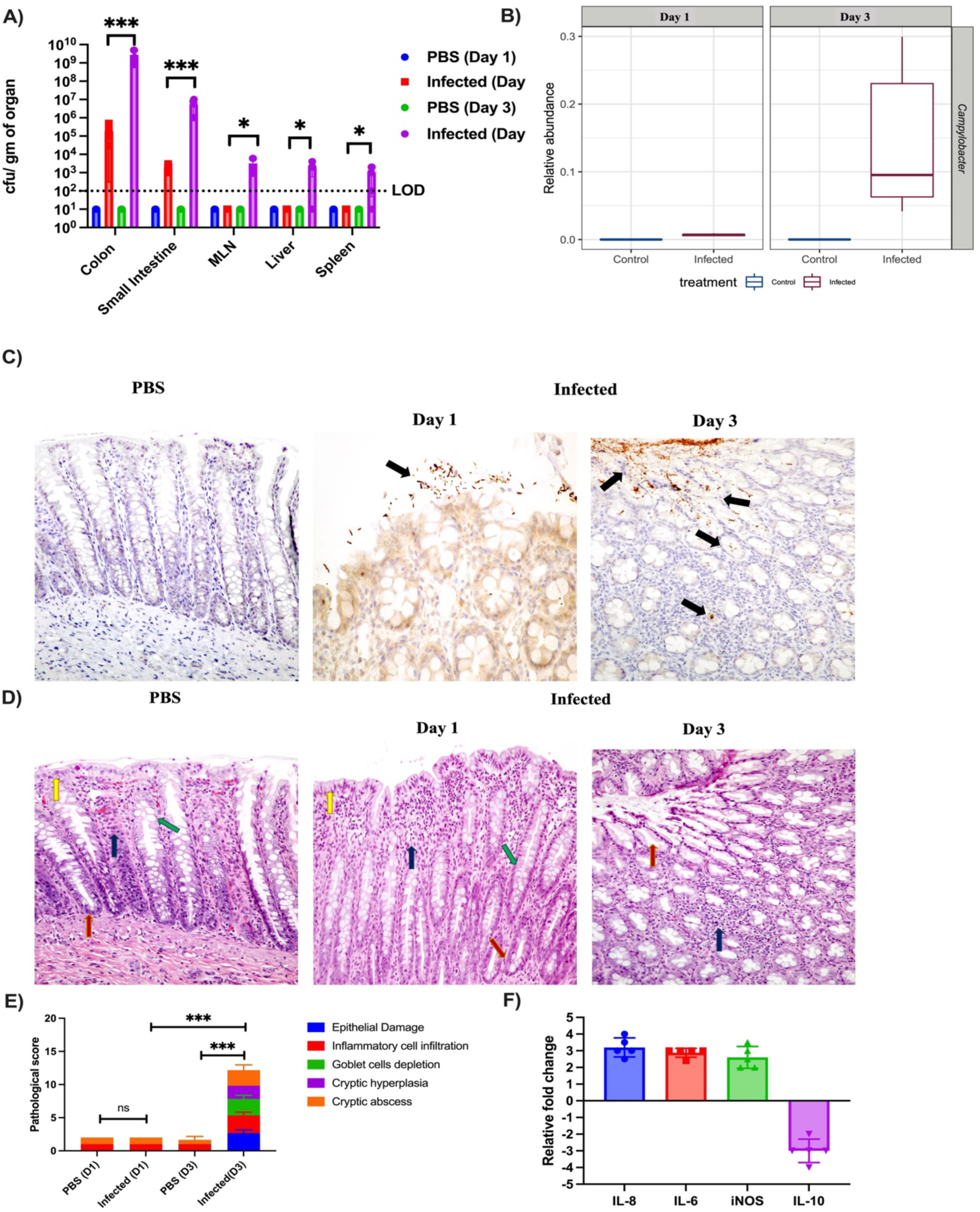
Characterization of ferret as a natural disease model for *C. jejuni* pathogenesis: A) *C. jejuni* 11168 WT loads were determined by CFU counts in different tissue samples Day 1 and Day 3 post infection with a dose of 10^9^ CFU/ml. (n=5). B) The relative abundance of *C. jejuni* was determined in colonic contents Day 1 and Day 3 post infection by 16Sr RNA sequencing. (PBS n=3, Infected n=6) C) Immunohistochemistry (IHC) with *C. jejuni* specific antibody was performed to determine *C. jejuni* localization in colon tissue Day1 and Day 3 post infection. Black arrows indicated localization of *C. jejuni* in infected colonic tissue. Representative images from two independent experiments. Original magnification, 40X. D) Histology of infected and PBS-shamed ferret colonic tissue. Representative hematoxylin and eosin (H&E) stained images on Day 1 and Day 3 post infection. Yellow arrow (epithelial cells), red arrows (intestinal crypts), green arrow (goblet cells), blue arrow (infiltrating leucocytes in lamina propria). Original magnification, 40X. E) *C. jejuni* mediated gastroenteritis as measured by histological score of infected and uninfected (PBS) colonic tissue Day 1 (D1) and Day 3 (D3) post infection (PBS n=3, Infected n=6). F) Relative fold changes of pro-inflammatory (IL-8, IL-6, iNOS) and anti-inflammatory (IL-10) cytokine genes determined by qRT-PCR analysis of infected colon tissue compared to that of the PBS control group (n=5). Changes in gene expression were determined by the 2^−ΔΔ*CT*^ method. All error bars show ± S.D. Statistical analysis was done by one-way AONOVA. *P < 0.05; **P < 0.01; ***P < 0.001; ****P < 0.000.

As *C. jejuni* levels were highest in the colon, we focused on the molecular mechanisms of colonization and inflammatory signs in this site. Analysis with anti-*Campylobacter* antibody identified sparse clusters of *C. jejuni* present within the colonic lumen area on day one; in contrast, day three sections demonstrated much greater levels of *C. jejuni* closely associated with colonic epithelial cells and detectable in crypts, like in human infection (**Fig 1C**)(3). To measure the intracellular burden of *C. jejuni* on day three after infection, we used a gentamicin protection assay previously described for *Shigella* and *Salmonella* in guinea pig and murine models, respectively (6). We incubated infected colon tissue with gentamicin, which kills extracellular bacteria, then determined colony counts of *C. jejuni* from the treated, washed tissue. After treatment, we observed approximately 10^4^ cfu/gram of tissue compared to approximately 10^8^ cfu/gram of tissue without gentamicin treatment. These data suggest that a portion of intestinal *C. jejuni* are present intracellularly (**FigS2A,B)**, which may lead to the systemic infection observed on day three. Through histopathological score analysis, we observed no significant changes in the inflammatory response of mock-infected and *C. jejuni-*infected animals on day one whereas by day three post-infection, we observed signs of moderate to severe colitis in infected colon tissue (**Fig 1D**). For example, we observed damage to the epithelial cell layer of infected tissue, infiltration of leukocytes in superficial lamina propria, and edema in infected colon tissue. This infected tissue also demonstrated evidence of crypt elongation into the gut lumen and a lower number of mucus-secreting goblet cells when compared to the mock-infected control group (**Fig 1E**).

Increased IL-6, IL-8 and iNOS levels prime leukocyte infiltration and tissue damage, and the anti-inflammatory IL-10 knockout mouse has been used to study pathogenicity of *C. jejuni* (7). To explore the inflammatory environment further, we measured mRNA levels from genes encoding these cytokines in colon tissue of mock-infected and *C. jejuni*-infected ferrets. Infection of ferrets with *C. jejuni* led to a two-to-three-fold increase in transcripts encoding pro-inflammatory IL-6, IL-8, and iNOS, and approximately three-fold decreased levels of transcripts from the anti-inflammatory IL-10 (**Fig 1F**). From the above analysis, we conclude that *C. jejuni* causes acute infection and inflammation in this model, resulting in cryptic hyperplasia in the colon on day three, like the acute stage of human infection (15).

During murine infection with *Citrobacterium rodentium,* cryptic hyperplasia is induced by virulence factors encoded on the locus of enterocyte effacement (LEE) (5). This induces tissue repair systems in the host, ultimately recruiting α-smooth muscle actin (α-SMA) myofibroblast cells and accumulation of undifferentiated proliferating epithelial cells (Ki67+ cells), which results in decreased numbers of goblet cells at the colon surface (5, 8). This type of tissue remodeling is associated with altered gut physiology that can support growth of the microbe. We therefore assessed whether these changes occur after *C. jejuni* infection of the ferret. Loss of goblet cells in infected tissue was observed by Alcian blue staining (**Fig 2A**). Immunohistochemistry (IHC) on colon samples from infected and mock-infected groups using antibodies for anti-α-SMA and anti-Ki67+ antigen demonstrated α-SMA myofibroblast cells recruitment (**Fig 2B**) and a significantly higher number of Ki67+ undifferentiated proliferated epithelial cells (**Figs 2C** **& D)** in the lamina propria of infected colons when compared to the mock-infected control group. We conclude that the environment of the ferret gut after *C. jejuni* infection is like that of the murine intestinal tract gut that supports *C. rodentium* growth.

**Figure 2:**
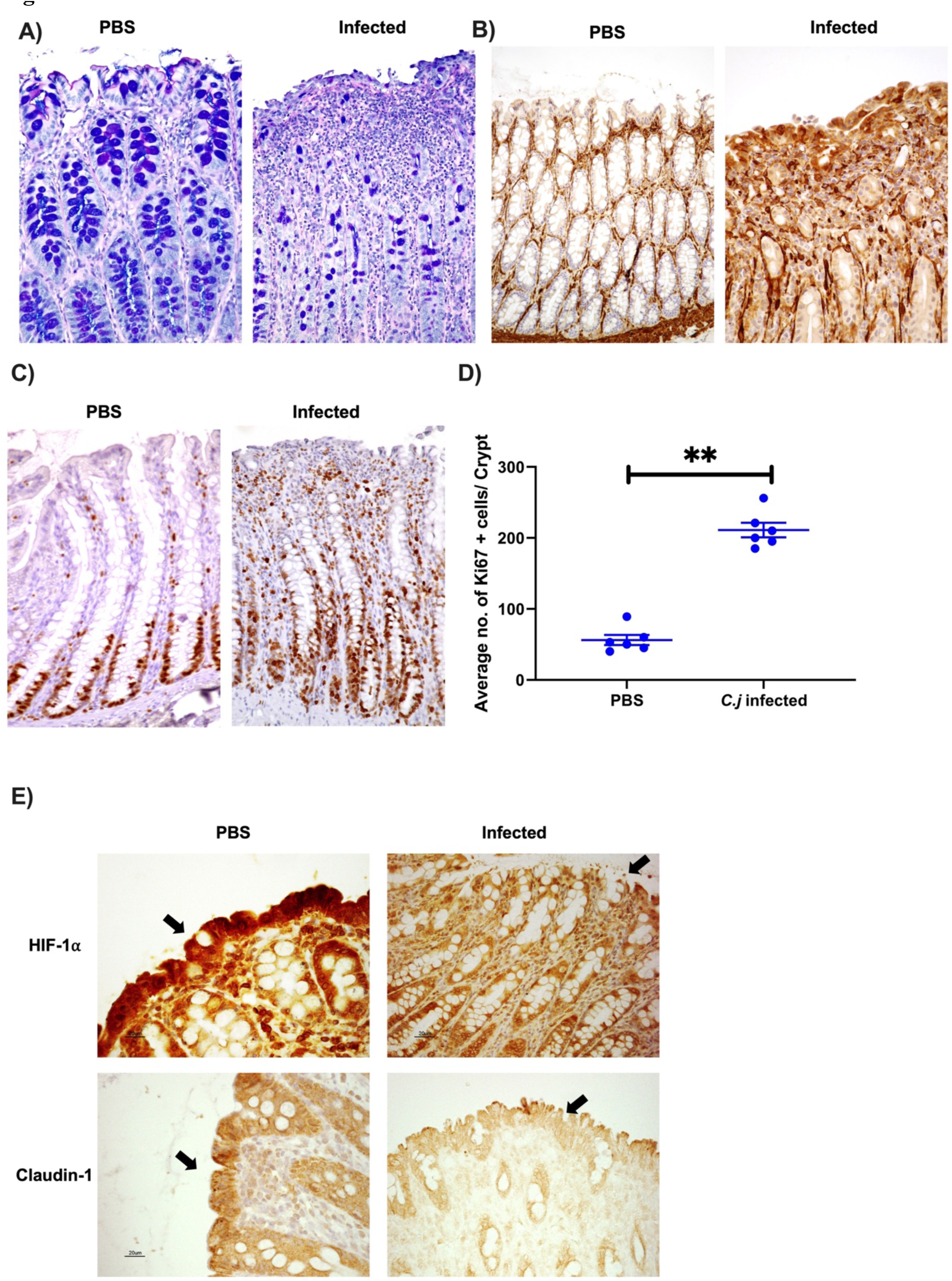
*C. jejuni* induced pathophysiological changes of colonic tissue during acute stage. A) Representative image of Alcian blue staining for goblet cells of colonic tissue Day 3 post infection. Original magnification, 40X. B) Representative IHC image of Day 3 post infection colonic tissue with α-Smooth Muscle Actin (SMA) specific antibody (to view tissue repair). C) Representative image of Ki67+ proliferated cell localization and D) count of Ki67+ cells/crypts in colonic tissue Day 3 post infection. Original magnification, 40X. All error bars show ± S.D. Statistical analysis was done by unpaired two-tailed t test. *P < 0.05; **P < 0.01; ***P < 0.001; ****P < 0.000.

Metabolism of microbially-derived butyrate by mature gut epithelial cells through beta-oxidation consumes oxygen, resulting in hypoxia within the gut lumen and stabilization of hypoxia inducible factor-alpha (HIF-1α) in colonocytes (9, 10). Undifferentiated proliferating cells, present mainly at the base of crypts lack ý-oxidation and use aerobic respiration that produces lactate from glucose(11). These cells do not absorb oxygen and as a result, the bottom crypt areas are more oxygenated than the top of the crypts(5). With the recruitment of undifferentiated, proliferating cells on the colon surface, we hypothesized that epithelial oxygenation is increased during the acute phase stage of *C. jejuni* infection.

To test our hypothesis, we performed IHC staining with anti-HIF-1α antibody from day three colon samples of both infected and uninfected animals. HIF-1α levels in colon sections from the infected group were lower than in those from mock-infected animals (**Fig 2E**), indicating a less hypoxic environment in the *C. jejuni*-infected animals, and suggesting epithelial oxygenation is increased during the acute stage of *C. jejuni* infection in the ferret. The tight junction protein claudin-1 is positively regulated by HIF-1 α (19), so we also analyzed tissue with anti-claudin 1 antibody. Consistent with the decreased level of HIF-1α in animals infected with *C. jejuni*, claudin 1 protein levels were also decreased in these animals compared to what we observed in mock-infected controls (**Fig 2E**). Loss of claudin-1 and subsequent disruption of tight junction integrity may play a role in the dissemination of *C. jejuni* in the ferret.

### Effects of *C. jejuni* infection on microbiota and metabolites in the ferret colon

The microbial composition in the colon contents of these five-to-six-week-old ferrets is made up predominantly of *Clostridia* sp., as determined by 16S rRNA gene sequencing **(Fig3A).** Dysbiosis and depletion of *Clostridium* after streptomycin treatment enables *Salmonella* Typhimurium to colonize the murine gut(12). Our data suggest that, in contrast, there is no negative effect of *Clostridium* on *C. jejuni* colonization in the ferret gut (12). Infected ferrets do not have a significantly altered overall microbiota (PERMANOVA, R2=0.129, Pr(>F) =0.097) **(Fig S3 A),** consistent with microbiota analysis of *C. jejuni*-infected humans (13). The low microbial diversity in the guts of 5-6 week old ferrets may help explain their susceptibility to *C. jejuni* infection.

By performing differential abundance analysis (DESeq2) of zero-radius operational taxonomic units (ZOTUs), we could identify differentially abundant taxa in colonic contents of the infected group from day three after infection. We observed elevated abundances of *Clostridium* sensu stricto 1 (ZOTU13), and (as expected) *Campylobacter* (ZOTU6) (log2 fold change = 25.13, p-adj=4.85e-16). This is reminiscent of *C. rodentium* infection in mice, which also resulted in greater *Clostridia* abundance (8). We also observed decreased abundance of *Enterococcus durans* (ZOTU12) (log2 fold change = -9.53, p-adj=5.40e-05) (**Fig 3B**). *Enterococcus durans* sp1 exhibits anti-inflammatory effects and reduced signs of DSS-induced colitis in mice (23, 24), suggesting perhaps that the reduced level of *E. durans* ZOTU12 abundance may contribute to induction of gastroenteritis in ferrets.

**Figure 3:**
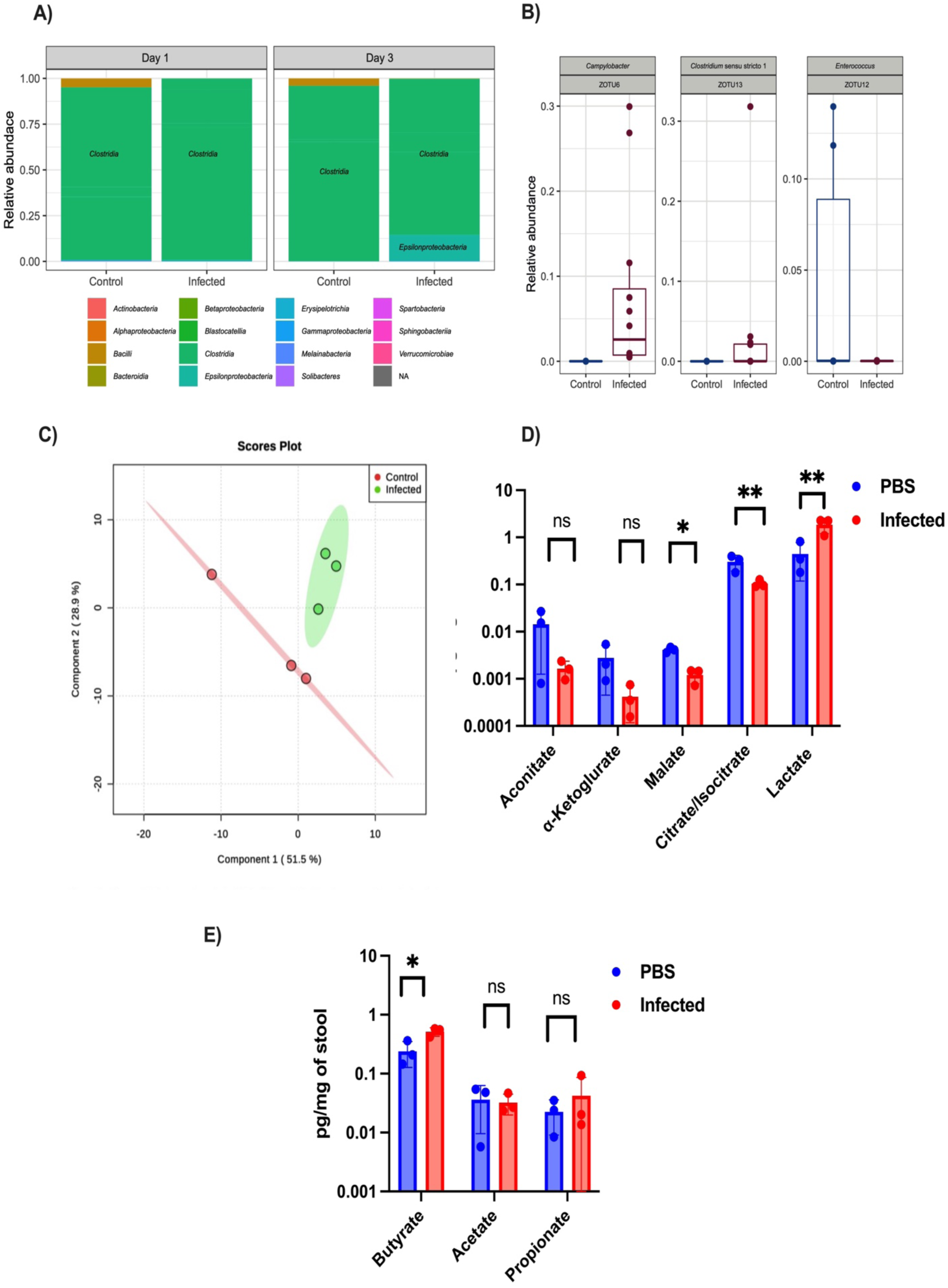
Changes of gut microbiota and metabolites after infection with *C. jejuni* in ferret. A) Microbial representation at class-level, based on 16s RNA sequencing, in cecal contents of infected and uninfected (PBS) ferrets Day1 and Day 3 post infection. Color coding for classes is shown below the chart. B) Differentially abundant taxa were found on Day 3 between infected and uninfected (PBS) ferrets using DESeq2. As expected, *Campylobacter* (ZOTU6) and *Clostridium* sensu stricto 1 (ZOTU13) members were significantly enriched in the infected group compared to the uninfected control group (log2 fold change = 8.93, p-adj=2.55e-06 and log2 fold change = 25.13, p-adj=4.85e-16, respectively). On the other hand, *Enterococcus durans* (ZOTU12) was the only one significantly decreased in the infection group (log2 fold change = -9.53, p-adj=5.40e-05). C) Difference of overall metabolites of colon contents between infected and uninfected control animals on Day 3 post infection by principal component analysis (PLSDA model). Each circle indicated overall metabolites of each animal, n=3. Concentrations of selected metabolites of colon contents from infected and uninfected animals on Day 3, such as D) different TCA cycles intermediates and E) short chain fatty acids, were determined by LC-MS mass spectrometry (n=3). Error bars represent SD. Statistical analysis was done by unpaired two-tailed t test. *P < 0.05; **P < 0.01; ***P < 0.001; ****P < 0.000.

Metabolites such as L-lactate, aspartate, malate, and formic acid, and others, which may be derived from the host or the microbiota, regulate growth and expression of virulence factors in various pathogenic bacteria (14–16). Less is understood about how gut metabolites influence *C. jejuni* growth during inflammation. One study determined that L-fucose, a component of mucin, induces *C. jejuni* growth and regulates colonization in a piglet model (17). Our histopathological analysis found that surface localized colonocytes were present after *C. jejuni*-induced colonic hyperplasia, which may influence the gut metabolites present during infection. To identify host-derived metabolites that might support *C. jejuni* growth during infection, we collected colon contents from uninfected and infected animals at 72 hours post-infection and subjected them to targeted mass spectrometry. Principal component analysis indicated significant difference in gut metabolite profiles between infected and mock-infected groups (**Fig 3C**). Of those metabolites found to be differently abundant, we focused on different known potential carbon sources for *C. jejuni*. Carbon sources that enable *C. jejuni* growth during infection are undefined. L-fucose is one of these, and the fucose transporter (*fucP*) is essential for colonization in a piglet model, although the fucose utilizing genes are not widely distributed among different strains (17). Studies carried out *in vitro* revealed that *C. jejuni* can use TCA cycle intermediates such as citrate, malate, α-ketoglutarate, and fumarate as carbon sources, and lactate use has also been reported (18, 19). We observed lactate levels elevated nearly 10-fold in colon contents of *C. jejuni-*infected ferrets while TCA cycle intermediates citrate, malate, and alpha ketoglutarate levels were significantly lower in samples from the infected group compared with those from the uninfected group (**Fig 3D**). Host-derived lactate serves as a sole nutrient source for several pathogens, and lactate utilizing genes are also essential for disease progression of *Salmonella* Typhimurium, *Neisseria* species and *H. influenzae* (14, 20, 21), Furthermore, short-chain fatty acids derived by the microbiota especially butyrate and propionate, reduce enteric colonization of pathogens such as *S. Typhimurium* and in and influence the inflammatory response in gut (12). Butyrate levels in colonic contents from infected ferrets were significantly higher than from the PBS control group, while other short chain fatty acid such as propionate and acetate were not significantly changed **(Fig3E**). Butyrate induces expression of the BumSR regulon, products of which are essential for commensal colonization of *C. jejuni* in the day-of-hatch chicken and in humans. (33). Based on the metabolite study, we sought to determine whether elevated lactate and butyrate in colonic contents of infected ferret could influence the growth of *C. jejuni* during inflammation.

### The lactate permease (*lctP*) operon is essential for *C. jejuni* expansion during ferret infection

We tested *C. jejuni* growth in mucin-containing minimal media supplemented with either 10mM butyrate or 10mM L-lactate and observed better growth with L-lactate in the media than with butyrate (**Fig 4A**). The genome of *C. jejuni* 11168 wild type contains four genes in an apparent operon *cj0076c-0075c-0074c-0073c*, where *cj0076c* encodes a lactate transporter (*lctP*) and *cj0075c-cj0074c-cj0073c* encode a non-flavin iron-sulfur containing three subunit membrane oxidoreductase which convert L-lactate to pyruvate. This operon is essential for L-lactate utilization *in vitro* (22). We hypothesized that lactate and *lctP* would be essential for growth during infection when lactate levels are elevated.

**Figure 4:**
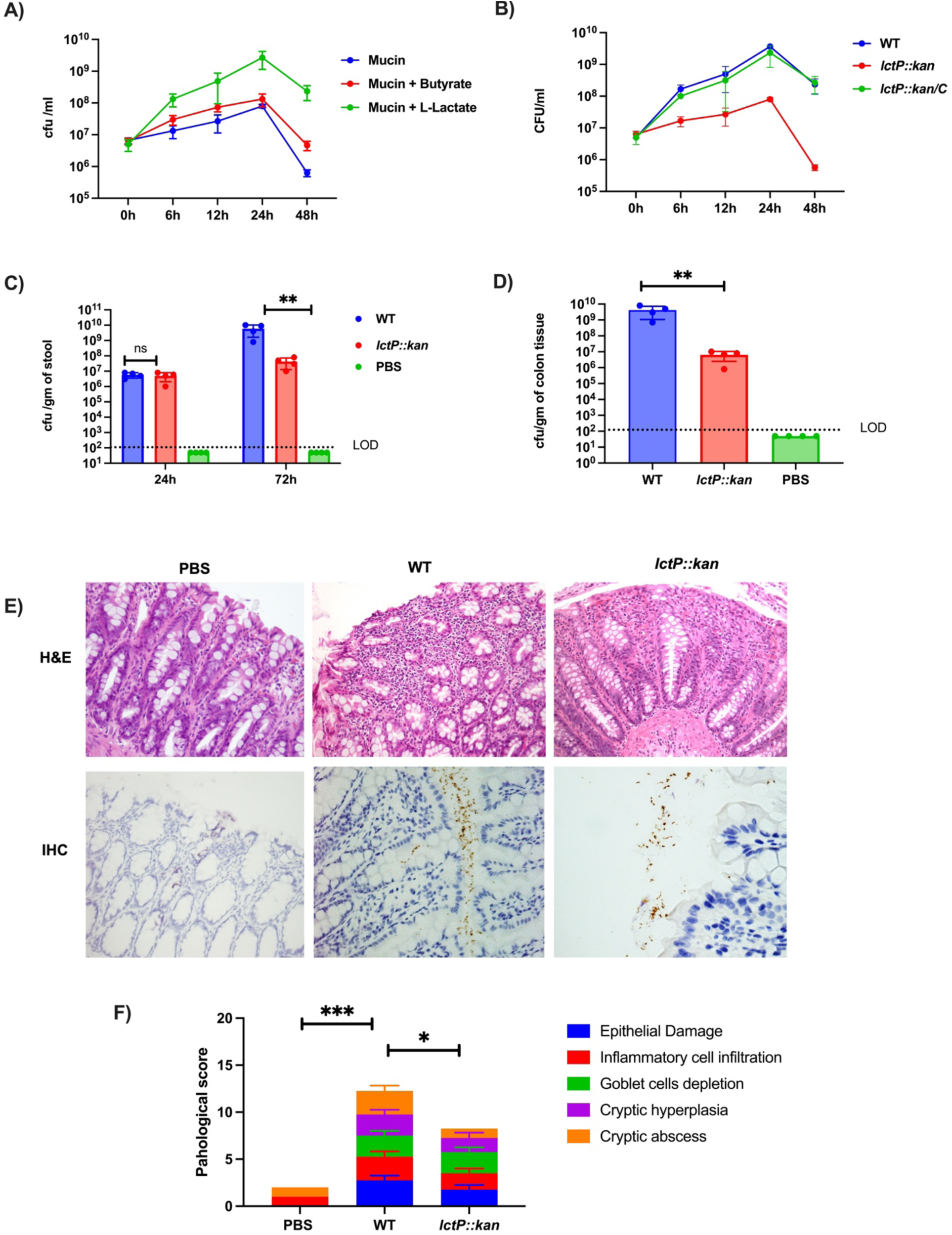
*lctP* operon is essential for pathogenesis of *C. jejuni* in Ferret. A) Growth of C. *jejuni* 11168 WT strain in minimal media containing mucin with and without butyrate (10mM) and L-lactate (10mM) over 48h was determined by CFU counts (n=3). B) WT, *lctP::kan* and complementary *lctP::kan/C* strains were grown in minimal media containing mucin with L-lactate (10mM); cfu/ml growth was determined over 48h (n=3). Bacterial comparative colonization was determined in 5-6 week old ferrets infected orally with either wild type or *lctP::kan* (dose of 10^9^ CFU/ml). Bacterial loads in C) stool and D) colonic tissue were determined by CFU counts at different time intervals. (PBS n=3 and infected group n=4). E) Representative images of H&E stained Day 3 colonic tissue (20X) and IHC with Day 3 colonic tissue using *C. jejuni* specific antibody (40X) (n =3). F) Histological scores of colonic tissue (PBS, WT and *lctP::kan* groups) were determined on Day 3 post infection (PBS n=3, Infected n=4). LOD indicates limit of detection. Error bars represent S.D. Statistical analysis was done by one-way AONOVA. *P < 0.05; **P < 0.01; ***P < 0.001; ****P < 0.000.

We constructed a mutation of *lctP* (*lctP::kan*) using a polar kanamycin-resistance cassette to knock down expression of the entire operon. RT-PCR of the wild-type locus using primers specific for the junctions between the open reading frames confirmed these genes are co-transcribed and that no detectable transcription of *cj0075c-cj0074c-cj0073c* is observed in the *lctP::kan* strain **(Fig S4A**). To complement this mutant, we amplified the entire operon and inserted it into the *lctP::kan* strain between the 16S rRNA and 23S rRNA genes using the prRNA-Hygro^R^ suicide vector (23). We measured growth of wild-type, *lctP::kan*, and the complemented *lctP::kan* strain in media containing 10 mM L-lactate. The wildtype and complemented *lctP::kan* strains grew similarly, whereas the *lctP::kan* mutant strain showed a significant growth defect achieving peak growth nearly 100-fold less than that of wild type and complemented strains (**Fig 4B**). Media in which wild type or *lctP::kan/C* were cultured retained low L-lactate while media in which the *lctP::kan* strain was cultured retained levels similar to the sterile culture medium. These data confirm that the *lctP* locus contributes to L-lactate uptake **(Fig S4B)**.

To examine whether lactate use is required by *C. jejuni* during infection, we infected ferrets with either wild-type *C. jejuni* or the *lctP::kan* mutant and measured fecal bacterial loads at 24- and 72-hours post-infection, and then in the colon of euthanized animals 72-hours post-infection. Fecal loads of wild-type and the *lctP::kan* were equivalent after 24-hours, but by 72-hours, wild-type *C. jejuni* was two orders of magnitude higher than the *lctP::kan* mutant (**Fig 4C**). Similarly, wild-type *C. jejuni* colonized the colonic tissue three orders of magnitude greater than the *lctP::kan* strain at 72 hours (**Fig 4D**). Supporting this result, immunohistochemistry demonstrated a higher number of adherent wild-type cells associated with the colonic epithelium with some cells colonizing deep into the colonic crypts. In contrast, very few *lctP::kan* cells were observed associated with the epithelial surface and they did not colonize within the crypts, remaining predominantly associated with the mucus (**Fig 4E**). In response to reduced infection, we also observed a diminished inflammatory score in colon tissue from *lctP::kan*-infected animals when compared to that of wild-type infected animals (although it was significantly higher than the PBS control group**) (Fig 4F)**. Overall, our ferret data suggest that the *lctP* operon is not essential for colonization by *C. jejuni* but is required for bacterial growth within the colon after colonization.

*C. jejuni* is a commensal microbe of chickens and establishes high loads after infection with limited inflammation that is not sufficient to clear the infection (5). Lactate is found in varying concentrations throughout the chicken gastrointestinal tract, typically decreasing as it progresses from the upper intestine to the lower intestine and ceca. Notably, in seven-day-old chickens, the average lactate concentration in the ceca is 4.2 ± 2.9 mmol/kg (24). We carried out competitive colonization assays between wild-type *C. jejuni* 11168 and the *lctP::kan* derivative in day-of-hatch chicks to assess whether lactate is a critical growth substrated for *C. jejuni*. Strains were mixed at a 1:1 concentration and inoculated into day-of-hatch chicks by oral gavage. We harvested cecal contents on day seven post-infection and determined the competitive index for each strain by measuring the number of kanamycin-resistant (*lctP::kan* mutant) and kanamycin-sensitive (wild-type) bacteria. There was no competitive disadvantage for the *lctP* mutant strain in chicks **(FigS5B)**, indicating that despite its availability, lactate uptake and utilization does not contribute to commensal colonization of chicks by *C. jejuni*.

### The *lctP* operon is required for adherence and invasion of human colonocytes

Metabolism of proliferating cells, such as the Ki67+ cells that are significantly elevated in infected ferrets, results in lactate production, which is the likely source of the elevated lactate we observe in infected animals. Similarly, human cancers have been found to produce and secrete lactate from metabolism of glucose, which has been termed the “Warburg effect” (11). To investigate whether host-derived lactate influences *lctP* expression, we introduced a plasmid containing a *lctP-gfp* promoter fusion into *C. jejuni* and infected HCT116 cells, a human colon carcinoma cell line. GFP levels were three-fold higher when bacteria were cultured with HCT116 cells compared to cultures lacking the HCT116 cells (**Fig 5A**). These data suggest that host-derived lactate produced by HCT116 cells induces expression of *lctP*.

**Figure 5:**
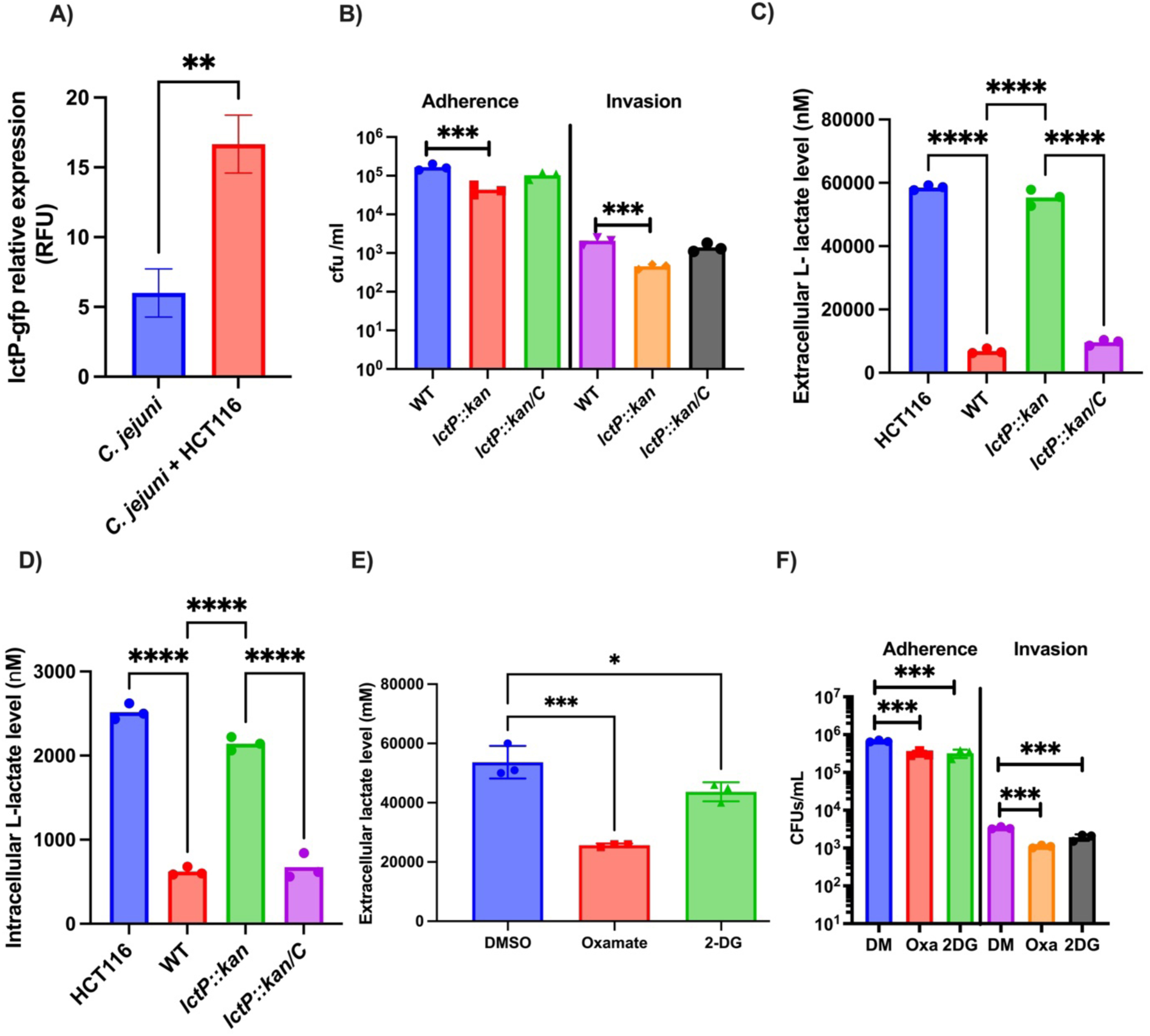
Host derived lactate influences the adherence and invasion of *C. jejuni* in vitro: A) Relative Fluorescence Units (RFU) of *C. jejuni* type strain carrying *lctP-gfp* fusion plasmid were measured with or without HCT116 cells after 1h of infection by spectrophometer. (n=3). Statistical analysis was done by unpaired two-tailed t test. B) Adherence and invasion ability of WT, *lctP::kan* and *lctP::kan/C* was determined after of 1 h of infection in HCT116 cells at an MOI was 10 (n=3). C) Extracellular and D) intracellular L-lactate levels in HCT116 cells were determined after 1 h of infection with WT, *lctP::kan* and *lctP::kan/C* by ELISA (n=3). E) Extracellular lactate level was determined after treatment with sodium oxamate (30mM) and 2-deoxy glucose (1mM) by ELISA (n=3). F) Adherence and invasion of WT was determined in sodium oxamate and 2-deoxyglucose and untreated HCT116 cells after 1 h of infection (n=3). Error bars represent SD. Statistical analysis was done by one-way AONOVA. *P < 0.05; **P < 0.01; ***P < 0.001; ****P < 0.000.

Because the *lctP* operon is needed for *C.jejuni* growth in the ferret colon, we explored the role of the *lctP* operon further by infecting HCT116 cells with either wild-type, *lctP::kan ,*or the *lctP::kan* mutant. After one hour of infection, we observed a small but statistically significant decrease in both adherence and invasion of the *lctP::kan* mutant when compared to wild-type and the complemented mutant (**Fig 5B**). This is consistent with what was observed for the wild-type and mutant during ferret infection, although the magnitude of the mutant defect during infection was greater than what we observe with cultured cells. To measure how the *lctP* operon contributes to lactate use from the host, we analyzed extracellular lactate levels in the culture media and intracellular lactate levels in cells after infection of the HCT116 cell line. Infection with either the wild-type or complemented *lctP::kan* strain resulted in lower lactate levels in culture media after one hour of infection, whereas we observed no significant change in lactate levels in the medium of cells cultured with the *lctP::kan* mutant. Similarly, we observed decreased intracellular lactate levels in cells infected with wild type and the complemented mutant, but not the *lctP::kan* mutant ((**Fig 5** **C, D).** We conclude that *C. jejuni* uses the LctP pathway for lactate uptake when extracellularly and intracellularly associated with host cells.

To further explore the role of lactate during *C. jejuni* infection of colonocytes, we treated HCT116 cells with two inhibitors i) sodium oxamate, a competitive inhibitor of lactate dehydrogenase and ii) 2-deoxyglucose, a non-competitive hexokinase inhibitor that reduces lactate production by inhibiting the glycolysis pathway (25, 26). Both inhibitors were confirmed to significantly reduce lactate levels when compared to untreated cells, without reducing cell viability (**Fig 5E**). We then infected treated and untreated cells with wild-type *C. jejuni* and measured bacterial adherence and invasion. We observed a slight, but statistically significant reduction in *C. jejuni* adherence and invasion in both treated groups when compared to the untreated controls (**Fig 5F**). These results suggest that host-derived lactate contributes to bacterial growth during adherence and invasion of *C. jejuni* to human colonocytes.

### LctR is a potential regulator of the *lctP* operon

Our data indicates that LctP-dependent lactate utilization promotes *C. jejuni* growth during infection, raising the question of whether *lctP* expression itself is regulated during infection. An earlier study examining transcriptome dynamics of *C. jejuni* demonstrated a reduction in *lctP* operon transcripts of over five-fold in cells undergoing transition from high (7.5%) to low (1.88%) oxygen levels (27). We used RT-qPCR to evaluate the expression of the *lctP* operon following a 2-hour incubation in mucin media containing lactate, conducted under both anaerobic and microaerobic conditions. Expression of *lctP* was reduced two to three-fold in the oxygen-limited condition when compared to the microaerobic condition (**Fig 6A**). These data suggest that *lctP* expression is influenced by environmental oxygen levels.

**Figure 6:**
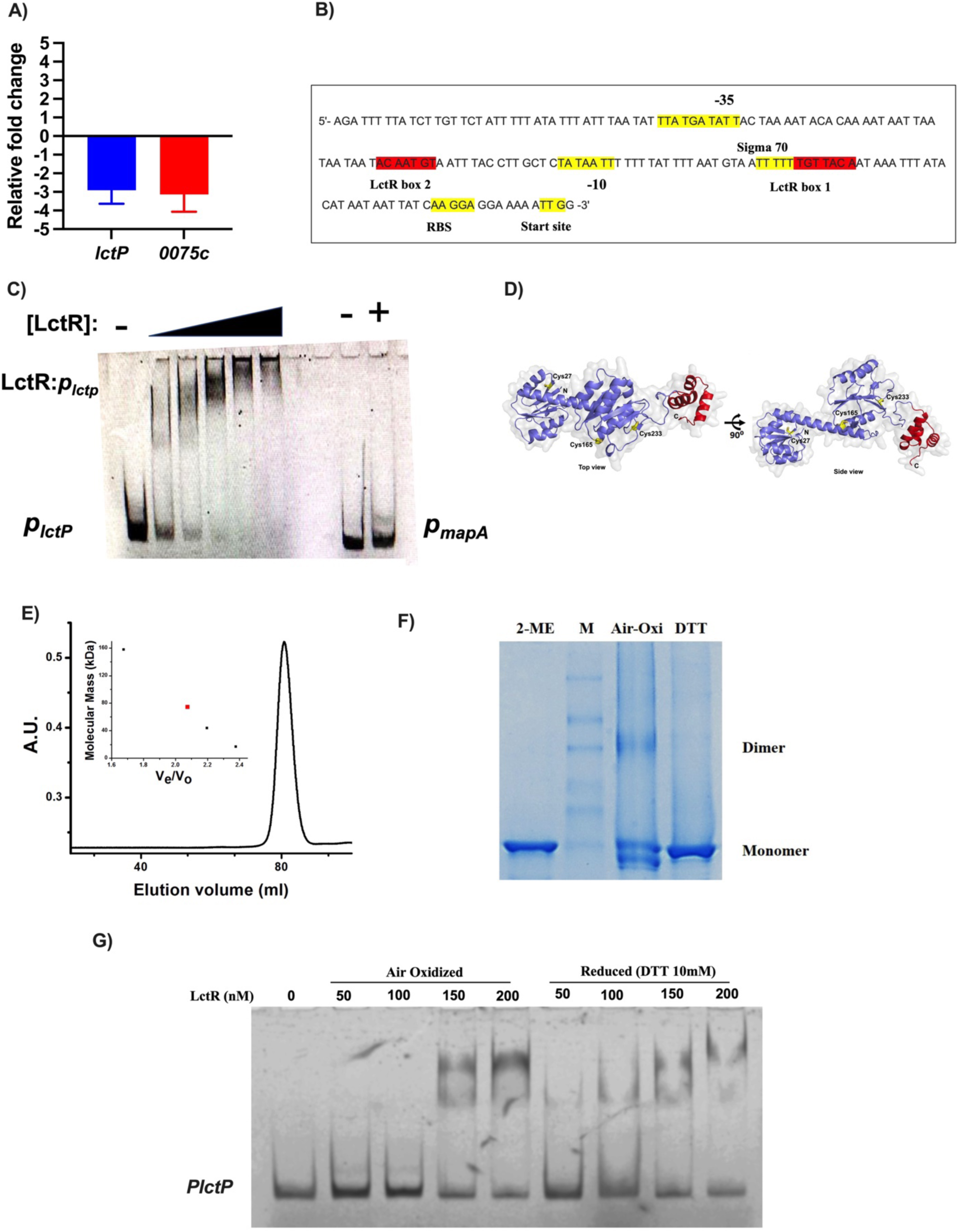
Redox switch protein LctR binds to the promoter of *lctP.* A) Transcription analysis of two *lctP* operon genes (*lctP* and *0075c*) at 2 h after growth of WT strain in minimal media containing mucin with lactate (10mM) by quantitative RT-PCR in anaerobic and microaerobic condition. Changes in gene expression of WT strain in anaerobic conditions compared with microaerobic conditions were determined by the 2^−ΔΔ*CT*^ method. Data are represented as the mean value from three independent experiments ± standard deviation (SD). B) Predicted promoter region of LctP (with important regions, such as –35, -10, and RBS, highlighted in yellow) and two probable LctR binding consensus sequences (5’-TGTTACA-3’, highlighted in red.) C) Determination of purified 6×His-LctR binding capacity to the promoter region of *lctP* (*P_lctP_*) by EMSA. Increasing amounts of purified 6×His-LctR were added to binding reaction mixtures containing approximately 0.25 nM either *P_lctP_* promoter fragment or the *mapA-ctsW* (*P_mapA_*) intergenic region as a nonspecific control. Results are representative of three biological replicates. D) A predicted structure of LctR by alphafold program. Three cysteine residues are highlighted in yellow and a predicted HTH domain is highlighted in red. E) Size exclusion chromatography of purified LctR. LctR eluted from Superdex column showed dimer form in normal condition. E) The thiol based redox state of LctR was analyzed by incubating with oxidizing (Air oxidation) and reducing (DTT) agent. Redox state of LctR was determined by 10% SDS page in non-reducing condition and Coomassie staining. Representative image of three biological replicates. F) Determination of LctR binding capacity to the *PlctP* promoter fragment in oxidized (Air oxidized) and reduced (DTT treated) condition by EMSA. Representative image of three biological replicates.

In other bacterial pathogens, *lctP* is regulated by oxygen and the ArcAB two-component system, which *C. jejuni* lacks (21, 28). During anaerobic growth of these microbes, the response regulator ArcA is phosphorylated by the sensor kinase, ArcB, and phosphorylated ArcA binds to the *lctP* promoter, repressing its expression. A redox-sensitive regulator in the related *Helicobacter pylori* is HP1021, which represses transcription of different genes such *gluP* and *fecA3* (among others) during growth in low oxygen (29). A homologous gene of *C. jejuni*, *cj1608*, encodes a product that binds to the *oriC* promoter via a high affinity consensus binding site (5’-TGTTACA-3’^)^ repressing replication of *C. jejuni* (a similar phenotype has been observed in *H. pylori*) (30). Two copies of this palindromic sequence are present upstream of the *C. jejuni lctP* operon at position - 25 to -85 relative to the transcription start site, which would place binding by CJ1608 within the potential RNA polymerase binding site (**Fig 6B**). Based on data presented in the next section we propose naming CJ1608 “***Lct***P ***R***egulator” (LctR).

We expressed and purified histidine-tagged LctR and used it in electrophoretic mobility shift assays with FAM-labeled *p_lctP_*. Increasing amounts of 6xHis-LctR gradually reduced the mobility of labeled *p_lctP_*, which was not observed for the non-specific binding control, a FAM-labeled region of the *mapA* gene (31) (**Fig 6C**). From this we concluded that 6xHis-CJ1608 specifically binds to *p_lctP_*.

Structural modeling of LctR with AlphaFold 2 indicated distinct N-terminal and C-terminal domains (29, 32, 33). The C-terminal domain includes a helix-turn-helix (HTH) motif (**Fig 6D**) common to DNA binding proteins, while the N-terminal domain includes three surface-exposed cysteine residues (Cys-27, Cys-165 and Cys-233; **Fig 6D**). The *H. pylori* HP1021 protein contains six cysteine residues, at least some of which control conformational changes of the protein in response to oxidation levels (29). Aligning sequences of HP1021 and LctR found that residue Cys-27 is conserved in the two proteins **(FigS6).**

Because our predicted model of LctR includes surface-exposed cysteines, we hypothesized that the protein may form cysteine-mediated oligomers, which may contribute to redox-sensitive changes in di-sulfide bond formation that could play a role in its activity. Size exclusion chromatography suggests that LctR exists as dimer in solution, with a molecular weight of 80 kDa (**Fig 6E**). Air-oxidized LctR migrates on non-reducing SDS-PAGE as multiple species, predominantly as a monomer but with evidence of dimer and even multimer formation (**Fig 6F**). The protein forms a dimer in prolonged storage conditions, as observed with gel filtration (**Fig 6F**). Incubating the protein with the reducing agent 10 mM DTT, we observed only a single species that migrates on non-denaturing SDS-PAGE consistent with its being a monomer, indicating that dimers and multimers are a consequence of thiol-oxidation. To further explore the impact of reducing or oxidizing LctR on its DNA binding ability, we tested four concentrations of LctR (50,100,150 and 200 nM), both air-oxidized after 30 minutes treatment with DTT (10mM). After exposure to reducing agent, LctR binding to *p_lctP_* was observed at lower concentrations of protein LctR (50nM and 100nM) than that observed for oxidized LctR (**Fig 6G**). Based on these results, we conclude that reduced LctR remains in a more stable, likely monomeric, conformation favored for DNA binding. Given the location of the putative LctR binding site in the *lctP* promoter, we suggest that LctR represses expression of the lactate utilization genes when bound in its reduced, monomeric form.

## Discussion

*C. jejuni* colonization results in different outcomes depending on the host: a commensal state, which occurs in the avian gut and results in little inflammation, and a pathogenic state in humans that leads to the development of intestinal inflammation and diarrheal disease (18). Because of these differences, the day-of-hatch chicken is a valuable natural host model for understanding molecular mechanisms of commensal colonization of *C. jejuni* in the avian gut but is unsuitable for gathering information about the disease state (18). In contrast, the weaning-aged ferret develops a self-limiting infection and intestinal disease similar to humans. Due to these similarities, we examined the changes to the intestinal environment during *C. jejuni* infection of ferrets to discover strategies the bacterium uses to support growth during acute infection.

We observed that *C. jejuni* infection led to moderate to severe gastroenteritis signs in ferrets. After infecting with 10^9^ bacteria, we observed that the *C. jejuni* population grew within two to three days, which was accompanied by cryptic hyperplasia and the accumulation of undifferentiated cells on the colon surface. During infection, we also observed instability of HIF-1α in colonocytes, which may indicate increased epithelial oxygenation. We detected no significant changes in the microbiota during *C. jejuni*-induced inflammation although we did detect elevated lactate, which we hypothesized to be a carbon source for *C. jejuni* growth during infection.

During gut homeostatic conditions, microbiota-derived short-chain fatty acids, especially butyrate, induce peroxisome proliferator–activated receptor γ (PPAR-γ) signaling in mature differentiated epithelial cells to maintain a state of physiological hypoxia in the lumen through oxygen consumption via β-oxidation (34, 35). Decreased oxygen availability in epithelial cells stabilizes HIF-1α which controls tight junction proteins, mucus production and antimicrobial peptides (36). The base of intestinal crypts is more oxygenated than the lumen because of the presence of undifferentiated amplifying epithelial cells which use anaerobic glycolysis and therefore consume less oxygen (35, 37, 38). The *Enterobacteriaceae* family of facultative anaerobic bacteria adopt different strategies to alter gut homoeostatic conditions for expansion in the gut (34, 35, 37). For example, *Salmonella* Typhimurium infection triggers neutrophil infiltration in the gut lumen, which causes depletion of *Clostridia* thereby increasing oxygen levels by altering the metabolism of mature colonocytes(34). *Salmonella* Typhimurium then also utilizes oxygen as an electron acceptor and lactate as a carbon source to grow in the gut lumen (28). In contrast, *Citrobacter rodentium* causes colon cryptic hyperplasia in mice through its type III secretion system (T3SS) which increases epithelial oxygenation by recruiting immature colonocytes (which lack β-oxidation metabolism) to the surface of the colon (34). During growth of *C. jejuni* in the ferret, we observed infiltration of neutrophils in infected colonic tissue, and prominence of *Clostridia* species and the metabolite butyrate which is quite different than *S. Typhimurium* infection in mice (37). We hypothesize that the strategy of *C. jejuni* expansion in ferrets during infection is more like *C. rodentium* in mice.

The obligate microaerophile *C. jejuni* requires low levels of oxygen (a partial oxygen tension (pO2) of 2–10%) for growth (39). Oxygen is an electron accepter for ATP production, growth, and motility of *C. jejuni*(39). We hypothesize that accumulation of undifferentiated immature colonic cells on the colonic surface during *C. jejuni* infection increases epithelial oxygenation. During inflammatory response to pathogens in the gut, epithelial oxygenation increases between 3 to 10% which indicates a microaerobic condition is created near epithelial cells (34, 35). This would create an appropriate environment for *C. jejuni* microaerobic respiration and growth in the gut (40). While the T3SS and its effectors in *Citrobacter rodentium* induce recruitment of proliferative undifferentiated epithelial cells during infection in mice, in the case of *C. jejuni,* the factors that induce colonic cell proliferation are not clear. *C. jejuni* secretes effector proteins, called Campylobacter invasion antigens (Cia), through a flagellum-associated mechanism, triggering signal transduction in host cells (5, 41–43). Bile salts induce synthesis of Cia proteins, and we detected elevated levels of primary bile salts and decreased leveled of one secondary bile salt, taurolithocholic acid (TLCA), which has an anti-inflammatory effect on LPS-induced macrophages, during the acute stage of infection **(Fig S3B)** (44, 45) . We hypothesize that during infection, elevated bile salts induce Cia proteins, leading to proliferating cells in the colon, which aids the growth of *C. jejuni*.

Nutrient resources, especially carbon sources, that support *C. jejuni* expansion in the inflamed gut are not well known. L-fucose is the only carbohydrate reported that induces growth and colonization (analyzed in a piglet model). Genes for catabolism of L-fucose are not well conserved among *C. jejuni* isolates from different sources (46). *C. jejuni* can metabolize some organic acids like citrate, butyrate, succinate, and lactate (47). Our targeted metabolite study found elevated lactate levels in colon contents; given that host derived L-lactate can serve as a major carbon source for *Salmonella* Typhimurium in a mouse model (28), we explored lactate as a potential growth substrate for *C. jejuni.* That the *C. jejuni lctP* operon mutant exhibited a colonization defect (Day 3) led us to conclude that lactate is a carbon source for *C. jejuni* during infection. Supporting this, the *lctP* locus (0076c-0073c) is present in 2138 out of the 2140 strains we queried from public collections such as *the Campylobacter* pubMLST website (http://pubmlst.org/campylobacter/), most of which were isolated from human patients **(TableS1)**. We did not focus on identifying the potential sources of elevated lactate levels in the gut (host- or microbiota-derived), however, we observed an accumulation of undifferentiated colonocytes, which produce lactate from glucose on the surface of the colon. In addition, we did not observe drastic changes in the microbiota that might account for new sources of lactate. Elevated lactate levels were similarly observed in *C. rodentium* infection of mice with the presence of undifferentiated colonocytes. Based on these data we hypothesize that elevated L-lactate in the ferret gut is also host-derived (48). Adherence and invasion of colorectal cancer epithelial cells (HCT116) by *C. jejuni* was reduced in the presence of two lactate inhibitors, sodium oxamate (LDH inhibitor) and 2-deoxyglucose (hexokinase inhibitor), suggesting that these traits, along with growth *in vivo*, are influenced by lactate levels. We concluded that host-derived lactate influences *C. jejuni* adherence and invasion.

In some pathogenic bacteria, thiol-based regulators play a vital role in responding to cellular stressors, including oxidative stresses, the production of virulence factors, and colonization (49, 50). In our study, we identified a potential thiol-based redox orphan regulator that we named LctR. This regulator bears similarity to an *H. pylori* redox regulator encoded by HP1021 that represses differential gene expression under anaerobic conditions (29). Our *in vitro* experiments suggest that expression of the *lctP* gene in *C. jejuni* relies on oxygen availability. Furthermore, we observed that the oxidized, dimeric form of LctR has reduced promoter binding capacity at the putative RNA polymerase binding site, compared to its reduced, monomeric form, which suggests a redox-dependent impact on transcription activation of *lctP*. Consequently, we hypothesize that reduced LctR represses *lctP* expression, such as under conditions of low oxygen availability. Our *in vivo* work demonstrates that expression of the *C. jejuni lctP* operon is crucial during the inflammatory stage, coinciding with evidence of rising epithelial oxygen levels in the gut. This observation allows us to draw a connection between the regulation of *lctP* by LctR, as demonstrated in our *in vitro* and *in vivo* study.

In conclusion, our findings provide a deeper understanding of how the host response to *C. jejuni* infection appears to contribute to growth of the microbe during infection.

## Materials and Methods

### Bacterial strains and media

All strains and plasmids used in the study are listed in Table S2. All *C. jejuni* strains were routinely grown under microaerobic conditions (85% N_2_,10% CO_2_, 5% O_2_) on Mueller-Hinton (MH) agar or in broth under necessary antibiotics at the following concentrations: hygromycine , 15 μg ml^−1^; kanamycin, 50 μg ml^−1^; trimethoprim (TMP), 5 μg ml^−1^. *C. jejuni* strains were stored in MH broth with 20% glycerol at −80°C.

### Construction of a *lctP*::*kan* insertion mutant and complementation

The *C. jejuni lctP*::*kan* mutant was constructed by inserting a kanamycin resistance cassette into the gene as previously described (51). Briefly, 500 bp of both upstream and downstream regions of the *lctP* locus was amplified by PCR with primers containing *Sma*I restriction sites. The amplified upstream and downstream regions were joined by PCR. The 1,000-bp amplified PCR product was cloned into the plasmid pGEM-T Easy. The *Campylobacter* kanamycin resistance cassette was excised from pILL600 using *Sma*I and ligated between the upstream and downstream cloned regions of the pGEM-T construct. The plasmid was electroporated into the wild type strain and grown on MH agar overnight at 37°C under microaerobic conditions. Cells were then plated and grown on MH agar containing kanamycin for 2 to 3 days, and successful integration of *lctP*::*kan* into the chromosome was confirmed by PCR. For complementation, we amplified and cloned the entire *lctP* operon between the 16S rRNA and 23S rRNA genes contained in the prRNA-Hygro^R^ suicide vector. This vector was electroporated into the *lctP::kan* mutant. All transformants were selected on MH blood agar plates supplemented with hygromycin B and confirmed by PCR.

### Growth analysis in MH broth and minimal media

WT, *lctP::kan*, and complemented (*lctP*::*kan/C*) *C. jejuni* strains were grown from freezer stocks on MH agar containing the desired antibiotics for 24 h under microaerobic condition. The strains were restreaked and grown for 16 h. The bacteria were suspended in MH broth and inoculated in 25 ml of MH broth containing the desired antibiotics to an initial optical density at 600 nm (OD_600_) of approximately 0.05. Bacterial numbers were measured at different times up to 48 h. To determine the growth of *C. jejuni*, strains were grown in MCLMAN minimal medium containing 0.5% partially purified gastric mucin and various carbon sources, including sodium pyruvate, sodium butyrate and L-lactate (28, 51), as noted. Growth was measured via absorbance at OD_600_ up to 48 h after inoculating at an initial OD_600_ of 0.05.

### Growth analysis in MH broth and minimal media

WT, *lctP::kan*, and complemented (*lctP*::*kan/C*) *C. jejuni* strains were grown from freezer stocks on MH agar containing the desired antibiotics for 24 h under microaerobic condition. The strains were restreaked and grown for 16 h. The bacteria were suspended in MH broth and inoculated in 25 ml of MH broth containing the desired antibiotics. The initial optical density at 600 nm (OD_600_) was approximately 0.05 and bacterial numbers were measured at different times up to 48 h. To determine the growth of C. jejuni strains in different concentrations of inorganic phosphate, strains were grown in 0.5% mucin containing MCLMAN minimal medium containing different carbon sources, including sodium pyruvate, sodium butyrate and L-lactate (37), as noted. Strains were washed twice with minimal medium and inoculated with 25 ml of minimal medium containing different phosphate concentrations. Growth was measured via absorbance at 600 nm (OD_600_) up to 48 h after inoculating at an initial OD_600_ of 0.05.

### Ferret infection

Pathogen free female domestic ferrets (Mustela putorius furor) at 5.5 – 6 weeks of age (generally ranging in weight from 240 – 468 grams) were purchased from Marshall BioResources (North Rose, NY) and housed in animal containment facilities at Michigan State University. Ferrets were restricted from food and water for 2-3 hours prior to inoculation. At the time of food/water restriction, ferrets were given 2 mg/kg Zantac (ranitidine hydrochloride) by intraperitoneal injection. Two to three hours after Zantac administration, ferrets were anesthetized with ketamine (25-40 mg/kg) by intramuscular injection. Once anesthetized, the ferrets were fed 10 ml of bacterial culture in sodium bicarbonate (5 ml of 1 x 10^9^/ml *C. jejuni* wild type or *lctP::kan* + 5 ml of 5% NaHCO3) by oral gavage. Uninfected control ferrets received a mixture of 5 ml PBS + 5 ml 5% NaHCO3. One hour after inoculation, the ferrets received 0.05-0.1 mg/kg lomotil orally, and again after 8 h. After infection, animals were monitored for development of diarrhea and dehydration. Weight and water consumption were recorded prior to infection and daily for up to 3 days post-infection. At 1- and 3-days post-infection, one group from each set of experiments were euthanized and feces and various tissues (liver, spleen, small intestine, and colon) were collected to determine the *C. jejuni* load. Samples were homogenized in sterile phosphate-buffered saline (PBS) and serial dilutions were plated on *Campylobacter* selective Mueller-Hinton (MH) agar containing 10% sheep’s blood, cefoperazone (40 μg/ml), cycloheximide (100 μg/ml), trimethoprim (10 μg/ml), and vancomycin (100 μg/ml). Plates were grown for 48 h under microaerobic conditions and colonies were counted.

For gentamicin assays, a 1- to 2-cm portion of uninfected and infected colons were cut open longitudinally and washed three times with PBS to remove luminal contents and then incubated in 1 ml of 1× PBS with 100 μg/ml gentamicin for 1 h at room temperature (6). The colon tissues were consequently washed three times with 1× PBS for 30 min with shaking. The tissue was then homogenized, and serial dilutions were plated on *Campylobacter* selective MH agar containing for enumeration of bacterial burden. The animal experiment protocol has been reviewed and approved by Michigan State University Institutional Animal Care and Use Committee (IACUC) with approval no. PROTO202100267.

### Histology and Immunohistochemistry

Colon tissues were fixed in 10% neutral-buffered formalin, followed by storage in 60% ethanol. Tissues were sectioned at 5 μm, mounted on frosted glass slides, and stained with hematoxylin and eosin (HE.) and Alcian blue (goblet cells). Blinded samples were numerically scored for sign of inflammation, such as epithelial damage, inflammatory cell infiltration, goblet cell depletion, cryptic hyperplasia, and cryptic abscess, as follows: 0, none; 1, minimal; 2, mild; 3, moderate; 4, marked; and 5 severe (52). To detect *C. jejuni* localization in colonic tissue by immunohistochemistry, a polyclonal anti-*C. jejuni* was used as the primary antibody. For the cell proliferation marker Ki67+ cells, Smooth muscle cells actin (-SMAα), hypoxia marker (HIF-1α) and Claudin-1 identified using anti-ferret Ki67+, -SMAα, HIF-1α and Claudin-1 antibody were used as a primary antibody. All histology and immunohistochemistry study were performed at Veterinary Diagnostic Laboratory facility, Michigan State University.

### 16S rRNA sequencing

Colon contents were collected, and the DNA was extracted from fecal samples using Qiagen Power Fecal kit per recommendation of the manufacturer. The 16S V4 region was amplified with the 515F-806R (53) primer pair and sequenced by the Illumina MiSeq sequencer (2×250bp paired end reads). 16S rRNA gene amplicon sequencing was performed at the Michigan State Genomics Core Research Support Facility. We processed the 16S rRNA gene using the UNOISE algorithm to perform error correction (denoising) of amplicon reads (53). Taxonomic annotations for ZOTU representative sequences were assigned in the QIIME 1.19 environment using SILVA v128 database (54, 55). All data and analysis are available in https://github.com/nejcstopno/gut_microbiome.

### SCFA Extraction

Frozen ferret colonic content samples (∼250 to ∼450 mg) were thawed on ice. Four volumes (i.e. 1.0 mL per 250 mg) of ice-cold 95% acetonitrile was added. Samples were briefly disrupted in an air-cooled Bead Bullet tissue homogenizer using 0.2 mm zirconium oxide beads and then shaken for 30 minutes at 4 degrees C. Samples were incubated overnight (∼16 hours) at –20 degrees C, then centrifuged for ten minutes at high speed (∼15,000 *xg*) at 4 degrees C to pellet solids. Supernatants were filtered through 0.2-micron nylon luer-lock syringe filters (Fisher Scientific).100 microliter aliquots were transferred to small volume inserts in autosampler vials with screw-top closures for LC-MS analysis.

For glycolysis intermediate metabolites extraction, samples on dry ice were combined with 800 microliters of ice-cold acetonitrile:methanol:acetone (8:1:1, v:v:v) and suitable stable isotope-labeled or synthetic internal standards, including D4-glycochenodeoxycholic acid and myristoyl phosphatidylcholine, for estimation of metabolite recovery and for relative quantitation across experimental groups. Samples were homogenized with zirconium oxide beads for 2 minutes in a Bullet Blender (Next Advance) at 4 degrees Celsius, incubated for 30 minutes at 4 degrees Celsius, then centrifuged for ten minutes at 10,000 × *g* at 4° C. Supernatants were filtered through 0.2-micron syringe filters (Fisher Scientific) and evaporated to dryness in a SpeedVac. Samples were resuspended in 95% acetonitrile for use in LC-MS analysis (56).

### Liquid chromatography-mass spectrometry

Short chain fatty acid identification utilized a Thermo model LTQ-Orbitrap Velos mass spectrometer operating in full scan negative ionization MS mode at 60,000 resolution over a mass range of 50-1000 m/z. The mass spectrometer was coupled to a Shimadzu Prominence HPLC system through an electrospray ionization source. Ten microliters of each SCFA extract was injected and separated on a Phenomenex HILIC LC column, 2.0 mm x 100 mm, 3.0 micron particle size, 100 Angstrom pore size, using a gradient of (A) water containing 50 mM ammonium formate, and (B) acetonitrile from 95% to 50% B over 3 minutes.A guard cartridge of matching chemistry was fitted to the analytical column.

Targeted metabolite identification used a Thermo model LTQ-Orbitrap Velos mass spectrometer operating in full scan negative ionization MS mode at 60,000 resolution with data-dependent high-energy collisional dissociation (HCD)-MS/MS over a mass range of 50 to 1,000 *m/z*. The mass spectrometer was coupled to a Shimadzu Prominence high-performance liquid chromatography (HPLC) system through an electrospray ionization source. Ten microliters of each metabolite extract were injected and separated on a Phenomenex HILIC LC column, 2.0 mm by 100 mm, 3.0-μm particle size, 100-Å pore size, using a gradient of water containing 50 mM ammonium formate and acetonitrile from 95% to 50% over 14 min as previously described (57). A guard cartridge of matching chemistry was fitted to the analytical column.

### Data Analysis

Short chain fatty acid species were identified on the basis of precursor ion accurate mass measurements and retention times compared against SCFA commercial standards (Sigma Aldrich), and quantification was performed against calibration curves. Compound identifications, isotope correction, chromatographic peak alignment, and peak area quantification were performed with MAVEN software). Multivariate statistical analysis was performed using MetaboAnalyst version 4.0 software (https://www.metaboanalyst.ca/) (57).

### Real-time PCR

Wild type strain *C. jejuni* 11168 was grown in two different conditions (anaerobic and microaerobic) *in vitro* in L-lactate containing minimal media and total RNA was isolated by the TRIzol method and RNeasy minikit (Qiagen). After digestion with TURBO Dnase treatment, cDNA was made by SuperScript III first-strand synthesis supermix (Invitrogen). Then real-time PCR was performed by the Syber green method (51). Expression of different targeted genes was measured by the 2^−ΔΔ*CT*^ method as previously described. Primer sequences or primers themselves for all genes examined in this study are listed in Table S2.

Inflammatory cytokines from infected and uninfected ferret colonic tissue were determined as previously described. Flash-frozen tissue was homogenized in the Trizol and total RNA was isolated by TRIzol method and RNeasy minikit (Qiagen) and digested with TURBO Dnase treatment. Then expression of cytokines IL-8, IL-6, NOS2 and IL-10 were determined by real time pcr (Syber green method) (58). Expression of different targeted genes was measured by the 2^−ΔΔ*CT*^ method as above described. Primers are listed in Table S3.

### *C. jejuni* adherence and invasion within intestinal epithelial cells

Wild-type *C. jejuni*, *lctP* mutant, and *lctP/C* cultures were grown on MH agar plates supplemented with TMP (5 μg/ml) and 10% defibrinated sheep blood for 48 h under microaerophilic conditions to use for invasion and adhesion assays. HCT116 cells were grown in Dulbecco modified Eagle medium (DMEM) supplemented with 10% fetal bovine serum (FBS), 1% L-glutamine, and 1% penicillin-streptomycin (Pen-Strep) to confluence, followed by seeding at 5 × 10^5^ cells per well and allowing to adhere overnight. Supernatant was aspirated off, and 500 μl of DMEM supplemented with 10% FBS and 1% l-glutamine was added to wells. One hundred microliters of the individual bacterial strains (at an OD_600_ = 0.5) were added to the appropriate wells and were briefly centrifuged at 3,000 rpm for 1 min. Wells were then incubated in aerobic conditions for 1 h at 37°C. For adhesion assays, the supernatant was aspirated, and the cells were washed three times in 1× PBS and then lysed by incubating in 0.5% Triton X-100 at room temperature for 5 min. Cells were then serially diluted in 1× PBS, plated on MH+TMP agar plates, and incubated at 37°C in microaerophilic conditions for 48 h to enumerate the bacteria. For invasion assays, 100 μl of DMEM containing 10% FBS and gentamicin sulfate to a final concentration of 100 μg/ml was added to the cells and allowed to incubate for 2 h at 37°C under microaerophilic conditions. After incubation, the supernatant was aspirated, and cells were washed three times in 1× PBS, serially diluted, and plated on MH+TMP to enumerate the bacteria inside the cells. For the sodium oxamate, 2-deoxyglucose, and CoCl_2_ treatments of HCT116 cells, 30mM, 1mM, and 100 μM substrates, respectively, were added to 5 × 10^5^ HCT116 cells one hour prior to bacterial infection. After the one hour chemical treatment, HCT116 cells were infected with *C. jejuni* and then continued through the adhesion and invasion assays described above. (59).

### *lctP* GFP expression assay

Wild-type and *lctP*-GFP mutant *C. jejuni* were grown on MH agar plates supplemented with TMP (5 μg/ml) and 10% defibrinated sheep blood for 48 h under microaerophilic conditions. HCT116 cells were grown in DMEM supplemented with 10% fetal bovine serum (FBS), 1% L-glutamine, and 1% penicillin-streptomycin (Pen-Strep) to confluence, followed by seeding at 5 × 10^5^ cells per well and allowing to adhere overnight in a black 24 well plate. The next day, the supernatant was aspirated and 500 μl of DMEM supplemented with 10% FBS and 1% L-glutamine was added to the wells. One hundred microliters of the individual bacterial strains (at an OD_600_ = 0.5) were added to the appropriate wells, briefly centrifuged at 3,000 rpm for 1 min, and allowed to incubate for 1 h at 37°C. One-hour post-infection, GFP fluorescence was quantified through excitation and emission wavelengths set to 488 and 520 nm, respectively, using a BioTek Synergy microplate reader. Transient expression of *lctP* was determined by quantifying GFP expression from *C. jejuni lctP*-GFP, subtracting out the background fluorescence of both the media and the HCT116 cells.

### Measurement of extracellular and intracellular lactate within intestinal epithelial cells

Wild-type *C. jejuni*, *lctP* mutant, and *lctP::kan/C* cultures were grown on MH agar plates supplemented with TMP (5 μg/ml) and 10% defibrinated sheep blood for 48 h under microaerophilic conditions to use for invasion and adhesion assays. HCT116 cells were grown in Dulbecco modified Eagle medium (DMEM) supplemented with 1% L-glutamine and 1% penicillin-streptomycin (Pen-Strep) in confluence, followed by seeding at 5 × 10^5^ cells per well and allowing to adhere overnight in a white welled 96 well plate. The following day, the supernatant was aspirated and 500 μl of DMEM supplemented with 10% FBS and 1% L-glutamine was added to the wells. One hundred microliters of the individual bacterial strains (at an OD_600_ = 0.5) were added to the appropriate wells, briefly centrifuged at 3,000 rpm for 1 min, and then incubated in aerobic conditions for 1 h at 37°C. The abundance of extracellular and intracellular lactate was determined using the Promega Lactate-Glo^TM^ Assay as follows. All reagents used for this assay were brought to room temperature before use. For extracellular quantification, spent media was first diluted 50-fold in 1x PBS. Next, the inactivation solution (0.6N HCl) was combined with diluted spent media in a white welled 96 well plate and mixed for 5 minutes. Afterwards, the neutralization solution (1M Tris base) was added and mixed for one minute, followed by adding the lactate detection reagent and mixing again for one minute. The plate was then incubated for one hour at room temperature before reading the luminescence using a BioTek Synergy microplate reader. For intracellular lactate quantification, spent media was aspirated and the cells were washed three times with ice-cold 1x PBS. After the final wash, 1x PBS was added alongside the inactivation solution. The plate was then mixed for 5 minutes. The neutralization solution was then added to the reaction and the plate was mixed again for one minute, followed by addition of the lactate detection reagent and a final mixing for one minute. The reaction was then incubated at room temperature for one hour, before luminescence was recorded using a BioTek Synergy microplate reader (25, 26).

### Protein purification

The gene *Cj1608 (lctR)* was amplified by PCR from *C. jejuni* 11168 genomic DNA using primers (Table S2). Fragments confirmed to have the full nucleotide sequence of *lctR* were excised using *Bam*HI and *Sal*I and ligated into the expression vector pET 28a (Novagen). Ligation products were transformed into T7 Express lysY/I^q^ Competent *E. coli* (NEB C3013). Transformants were confirmed using a restriction digest of purified plasmids and sequencing. Induction of 6×His-LctR using 1 mM IPTG (isopropyl-β-d-thiogalactopyranoside) was confirmed for six constructs by SDS-PAGE and Coomassie staining. One strain was used for protein purification where a 1-liter culture was grown to the mid-log phase and induced for 3 h at 37°C using 1 mM IPTG. Cells were pelleted at 8,000 rpm for 20 min at 4°C and resuspended in 20 ml lysis buffer (pH 8.0) (50 mM NaH_2_PO_4_, 300 mM NaCl, 10 mM imidazole). Concentrations of 6×His-LctR were determined using a bicinchoninic acid (BCA) protein assay (Pierce), and aliquots of protein were stored at −80°C (31).

### Thiol redox state of LctR *in vitro*

To Investigate the redox state of LctR *in vitro*, LctR (2.72 μM) were diluted in Tris buffer and treated with 10 mM DTT. For reduction state of LctR, DTT treatment was done for 30 min at room temperature. The samples were heated at 95°C in non-reducing loading buffer and separated by electrophoresis on a 10% SDS-PAGE gel and coomassie blue staining was done.

### Electrophoretic mobility shift assays

A FAM labeled DNA probe specific for the *lctP* promoter region (at position -25 to-85 bp upstream of terminal transcription start site) and a FAM labeled nonspecific probe for the *mapA*-*ctsW* intergenic region (at position -196 to -7 bp upstream of transcription start site) were generated by PCR of *C. jejuni* 11168 genomic DNA and purified using a PCR purification kit (Qiagen) **(Table S2)**. FAM labeled specific (*lctR* promoter) and nonspecific probes at approximately 0.25 nM, DNA binding buffer (10 mM Tris, pH 7.5, 50 mM KCl, 1 mM EDTA, 5% glycerol), 25 ng/µl poly(dA-dT), and 100 µg/ml BSA were mixed to make the binding reaction mixture. This mixture was incubated for 5 min at room temperature before adding purified 6×His-LctR at different concentrations (0-800 nM) of protein for a total reaction volume of 20 µl. These reaction mixtures—and a control reaction mixture without 6×His-LctR—were incubated for 15 min at room temperature and subsequently resolved on a 5% polyacrylamide native Tris-glycine gel at 4°C. Gels were imaged using a Typhoon phosphoimager (43, 45).

### Size exclusion chromatography

The oligomeric state of LctR was analyzed by size exclusion chromatography on a BIORAD BioLogic DuoFlow chromatographic system using a HiLoad/GE-Healthcare Superdex 200 pg column (16 × 600 mm; bed volume ∼120 ml) pre-equilibrated with lysis buffer (50 mM Tris, pH 7.0, 150 mM NaCl) running at 298 K with a flow rate of 1.0 ml min^−1^. The elution profile was determined by monitoring the absorbance at 280 nm. γ-globulin (158 kDa), ovalbumin (44 kDa), and myoglobin (17 kDa) were used as molecular mass standards (60).

## Supporting information

LctP distribution among isolates

Supplemental Tables, Primers

## Acknowledgments

This work was supported in part by NIH award AI111192 and the Michigan State Rudolph Hugh Endowment (to V.J.D.) and AI166535 (to J.G.J.). S.C. was supported by NIH award CHE2203472 and NIH award GM128959 (to R.P. Hausinger and J. Hu), N.S. was supported by the Michigan State University Plant Resilience Institute. S.M.C was a trainee of the SEC Emerging Scholars Program.

## Author contributions

R.S. and V.J.D. designed research; R.S., R.M.L., S.M.C., S.C., performed research; N.S., M.T., J.J., R.S., V.J.D analyzed data, R.S., V.J.D. wrote the manuscript, R.M.L, S.M.C, J.J edited the manuscript.

## Supplementary Figures

**Figure S1:**
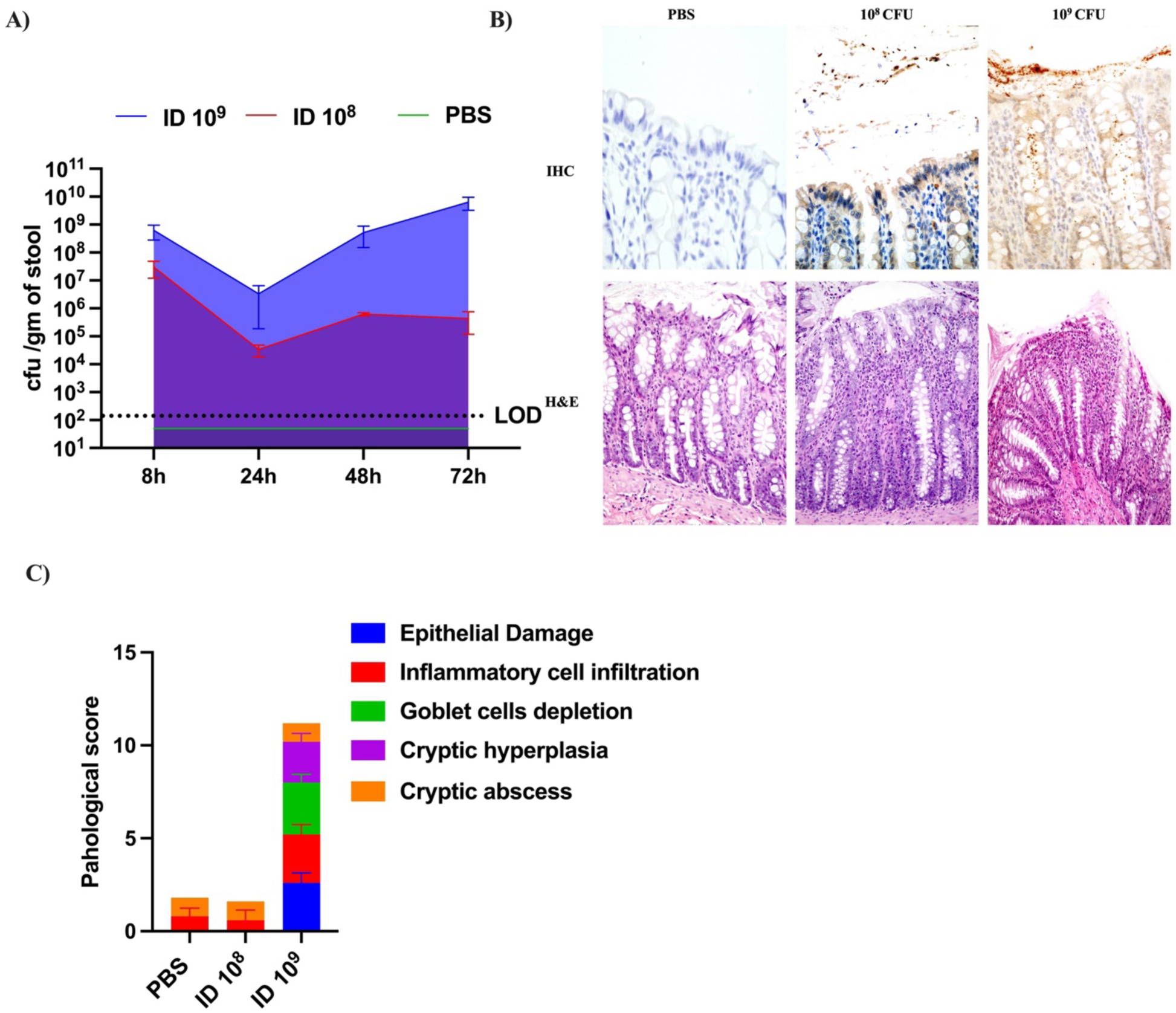
Dose dependent response of *C. jejuni* infection in ferret. 5-6 weeks old ferrets were infected with two different doses (10^8^&10^9^CFU/ml) and A) *C. jejuni* loads in the feces were determined by CFUs after different time intervals, (n=5 of each group). B) On Day 3 post infection immunohistochemistry with *C.jejuni* specific antibody was performed to determine localization of *C. jejuni and* Hematoxylin and Eosin (H&E) staining was used to measure *C. jejuni* mediated gastroenteritis. C) *C. jejuni* mediated gastroenteritis was measured in colon tissue of infected and uninfected (PBS) by histological score of infected colonic tissue after Day 3 post infection.

**Figure S2:**
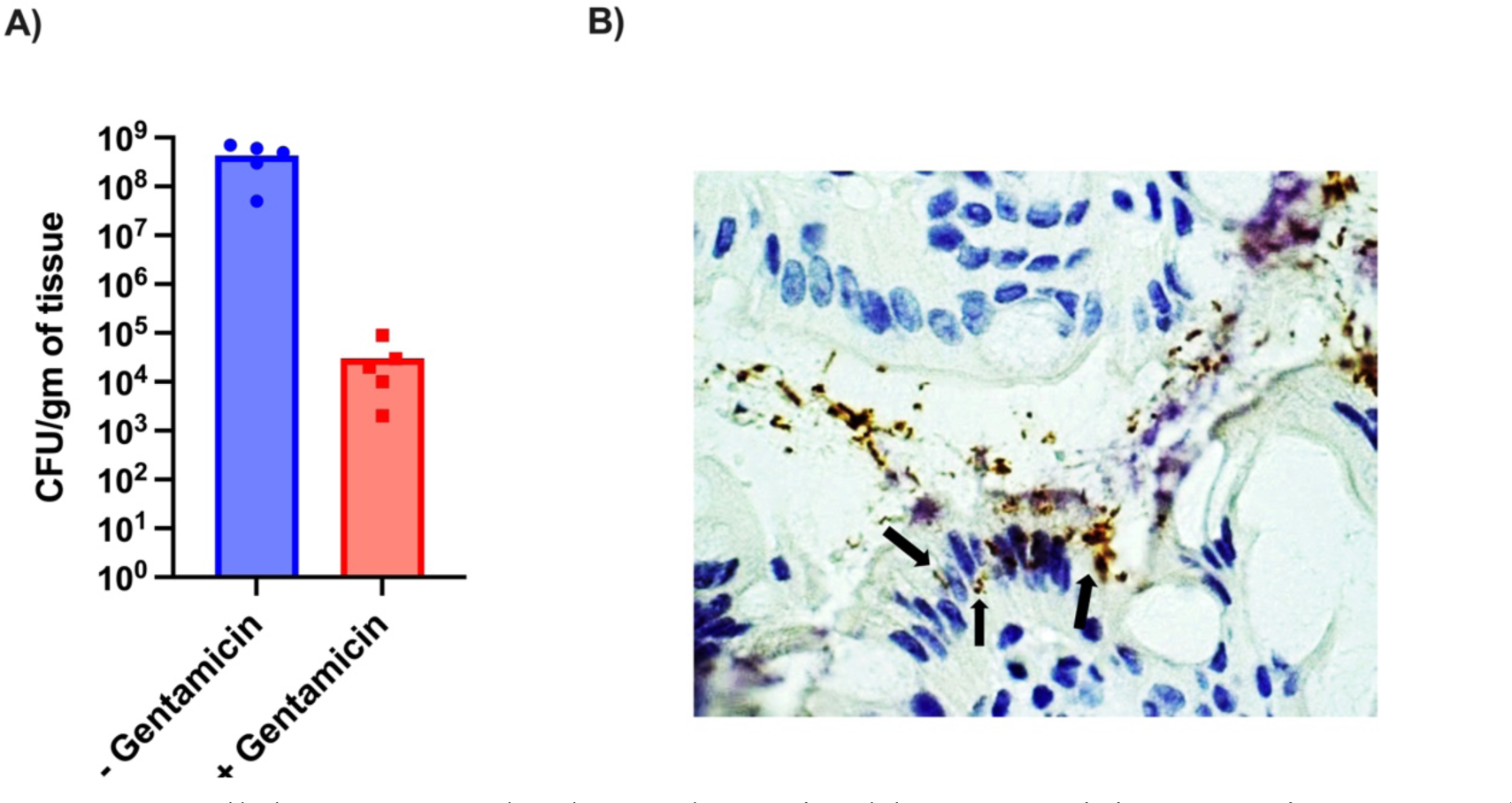
Intracellular *C. jejuni* load was determined by gentamicin protection assay. A) *C. jejuni* burden were determined by CFU count from Infected ferret colonic tissues after treatment with and w/o gentamicin (100ug/ml) for 1 h. n=5. B) Immunohistochemistry with *C.jejuni* specific antibody was done to determine invasives of *C. jejuni* in colon tissue on Day 3 post infection. Localization of invasive *C. jejuni* were determined by immunohistochemistry. The black arrows point to intracellular.

**Fig S3.**
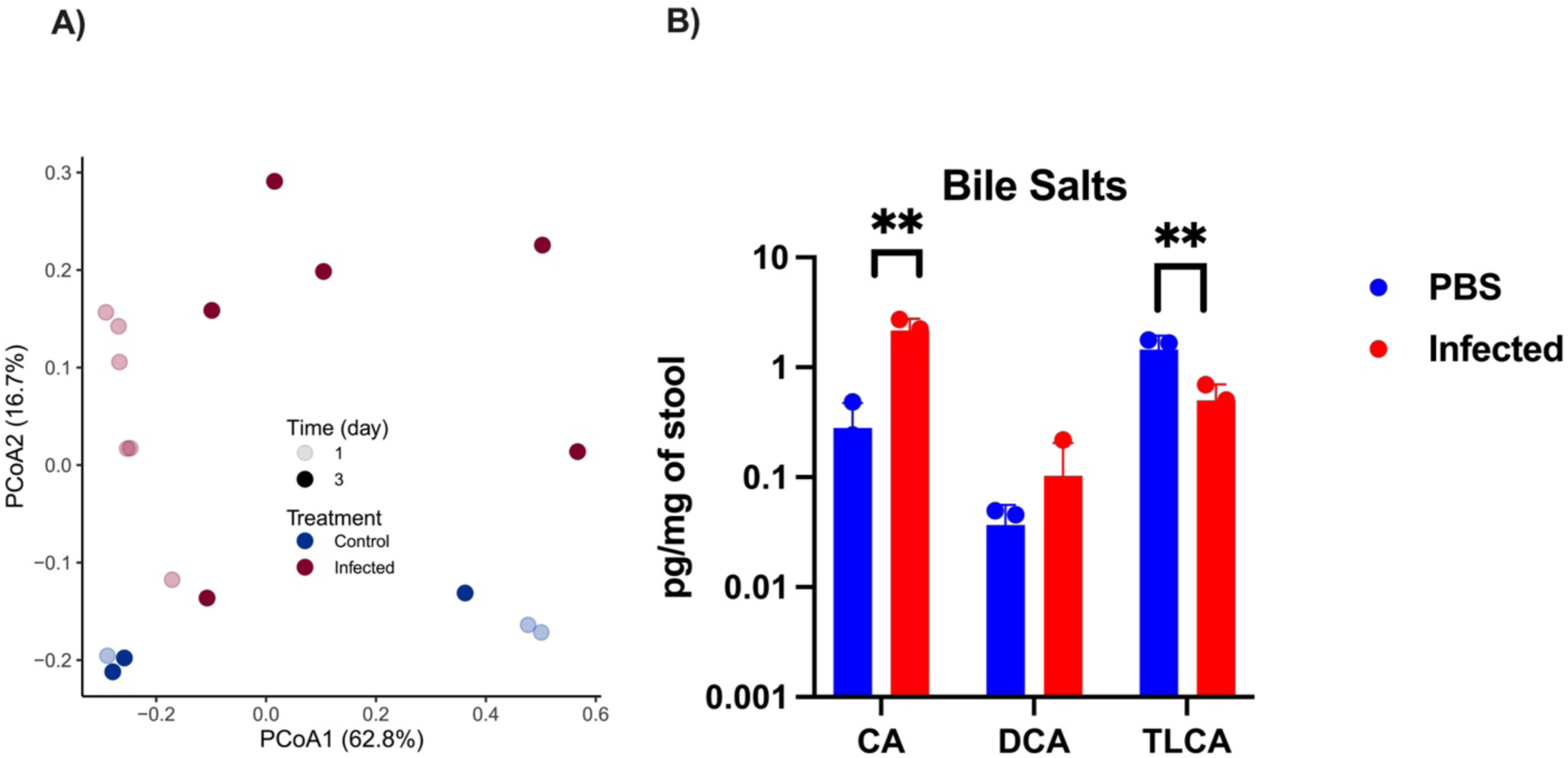
A) Community structure is not affected by *C. jejuni* infection (PERMANAOVA, R2=0.129, Pr(>F) =0.097) or time (PERMANOVA, R2=0.088, Pr(>F) =0.18). Time is shown as color gradient with lighter color representing day 1 and darker day 3 samples. (Control and infection treatments are represented by blue and red color, respectively.) B) Primary and secondary bile salts concentrations of colon contents of infected and uninfected on Day 3 were determined by LC-MS mass spectrometry (n=3). Error bars represent SD. Statistical analysis was done by unpaired two-tailed t test. *P < 0.05; **P < 0.01; ***P < 0.001; ****P < 0.000.

**Figure S4:**
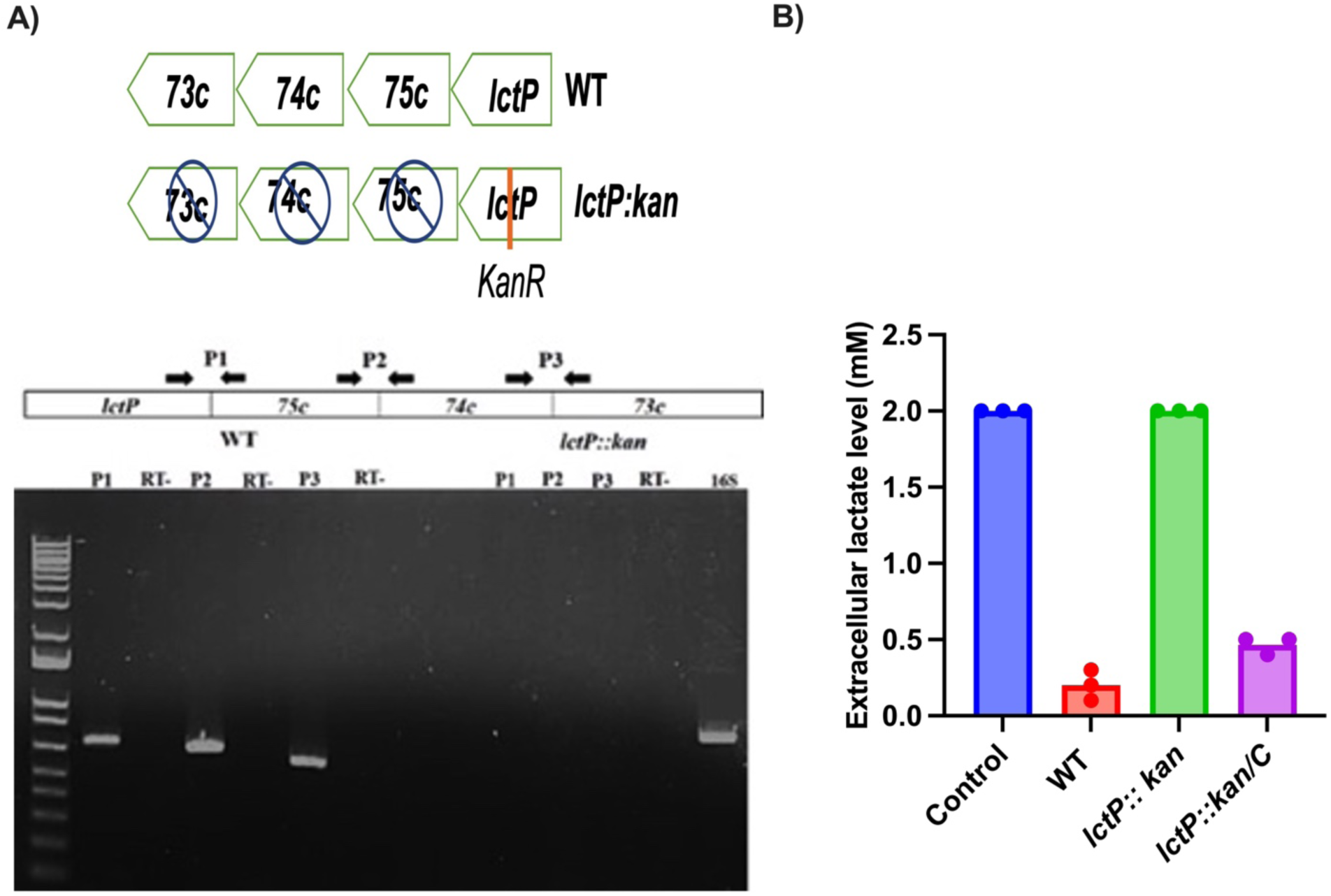
A) Schematic diagram and characterization of *lctP* operon containing complementary four genes (*lctP*, 0075c,0074c and 0073c) and RT PCR was performed to determine that these four genes are co transcribed. Three intergenic primer sets (P1, P2, and P3) were designed to amplify transcripts crossing the gene boundaries. The product sizes in base pairs were correct as predicted in all cases. RT-PCR with total RNAs were performed from both the wild type (left) and *lctP*::*kan* mutant (right) using the designed three primer sets. RT-, negative control; 16S, positive control. B) WT, *lctP::kan* and complementary *lctP::kan/C* were grown for 2h in minimal media containing 2mM L-lactate and Extracellular L-lactate concentration in media was different determined b**y** ELISA (n=3). Error bars represent SD. Statistical analysis was done by One Way Anova. *P < 0.05; **P < 0.01; ***P < 0.001; ****P < 0.000.

**Figure S5:**
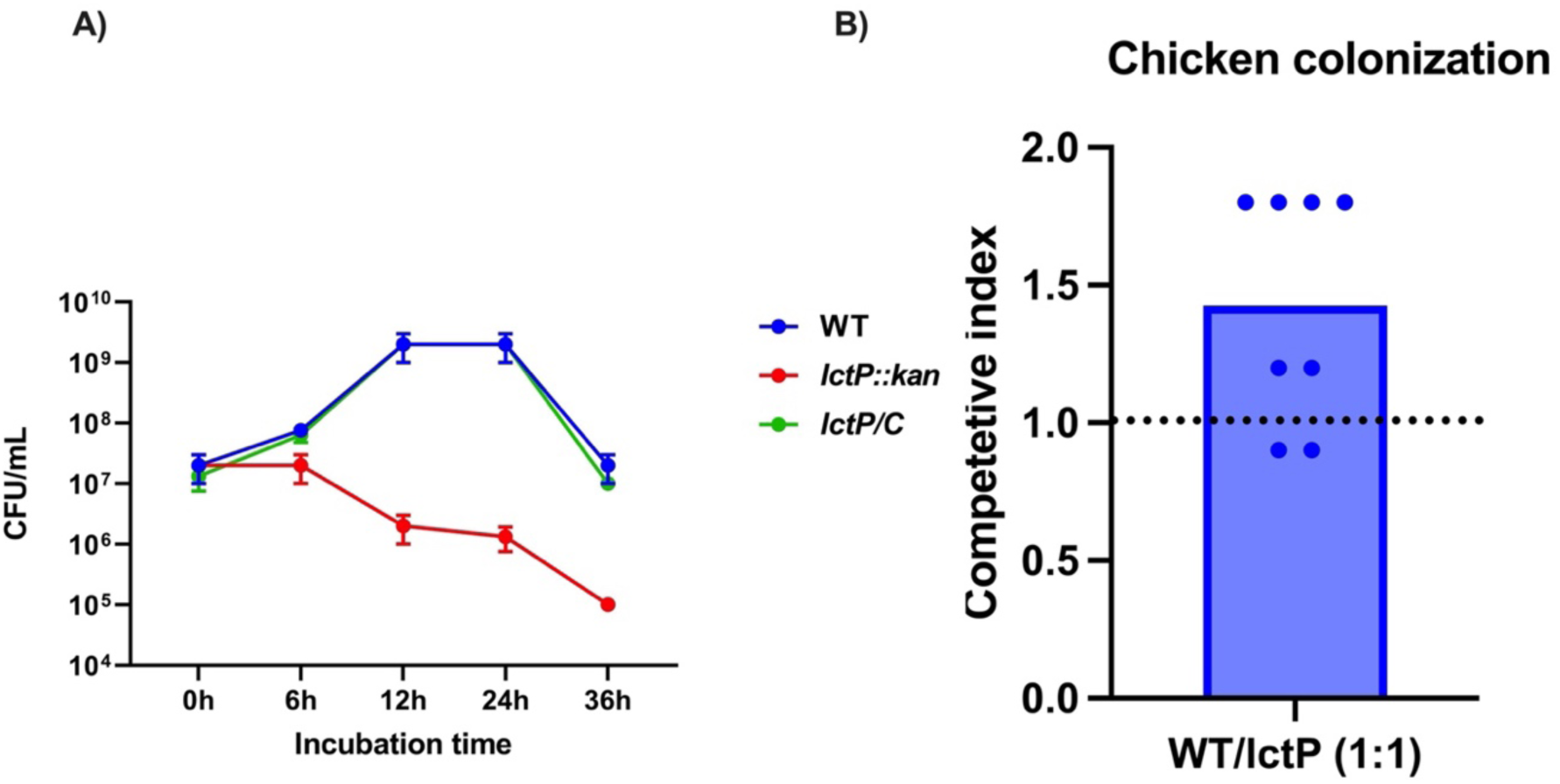
A) Growth of WT, *lctP::kan* and complementary *lctP::kan/C* strains were grown in minimal media with L-lactate (10mM) and cfu/ml were determined over 48h (n=3). B) In competition analysis between WT and the *lctP::*kan mutant, day-of-hatch chicks were infected with equal ratio (1:1) with WT and *lctP::kan* mutant strain. On day 7, the ratio of *lctP::kan* mutant to WT strain was determined and is represented as a competitive index. Statistical analysis was performed using a one-simple *t* test against a hypothetical value of 1 (*n* = 8 for each group). *, *P* < 0.05; **, *P* < 0.005.

**Figure S6.**
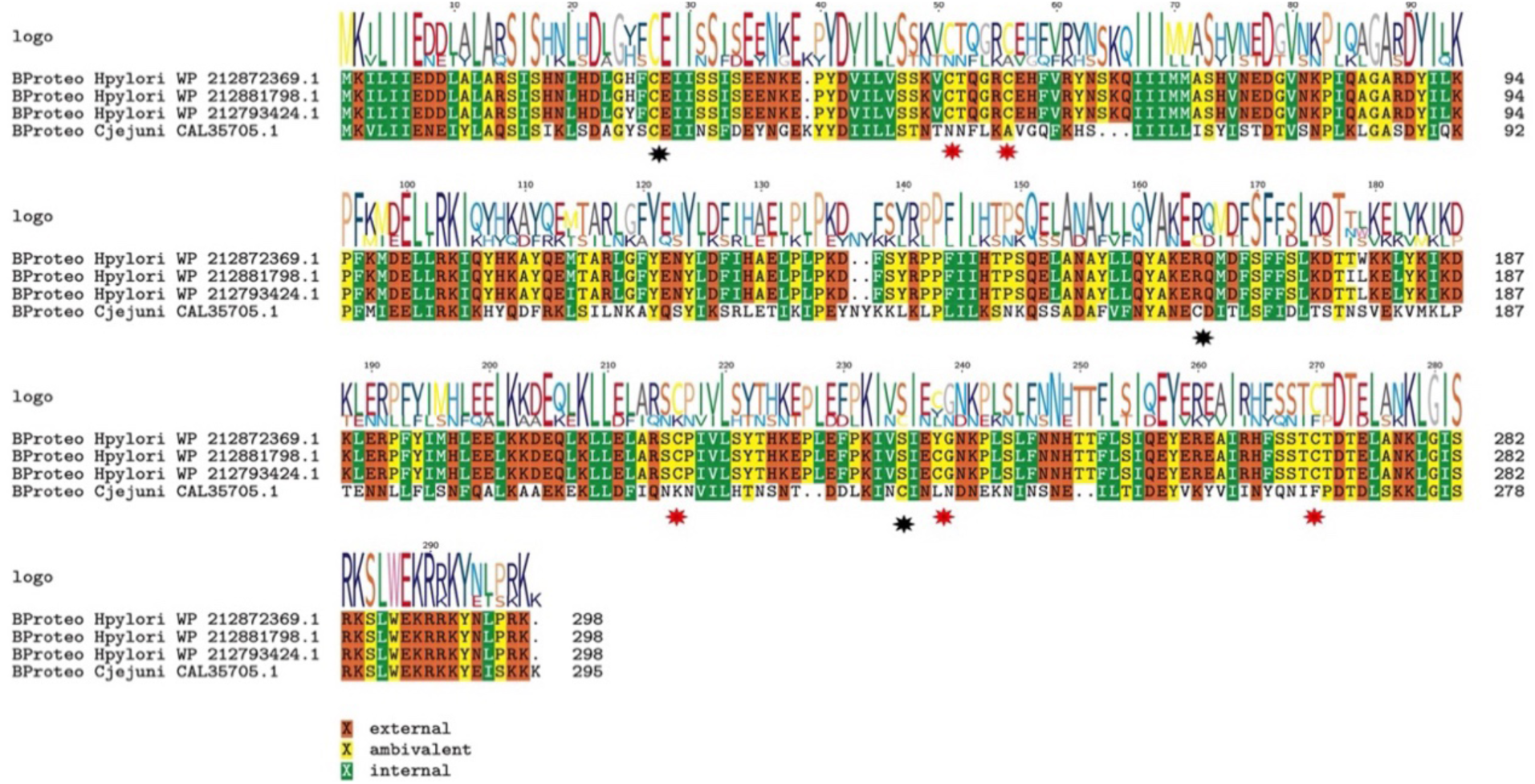
A) Amino acid sequence alignment of the LctR and response regulator HP1021 of *Helicobacter pylori* by MolEvolvR. Black stars indicate cysteine residue in C.jj 1608 (LctR) and red stars indicate cysteine residue in *H. pylori* HP1021.

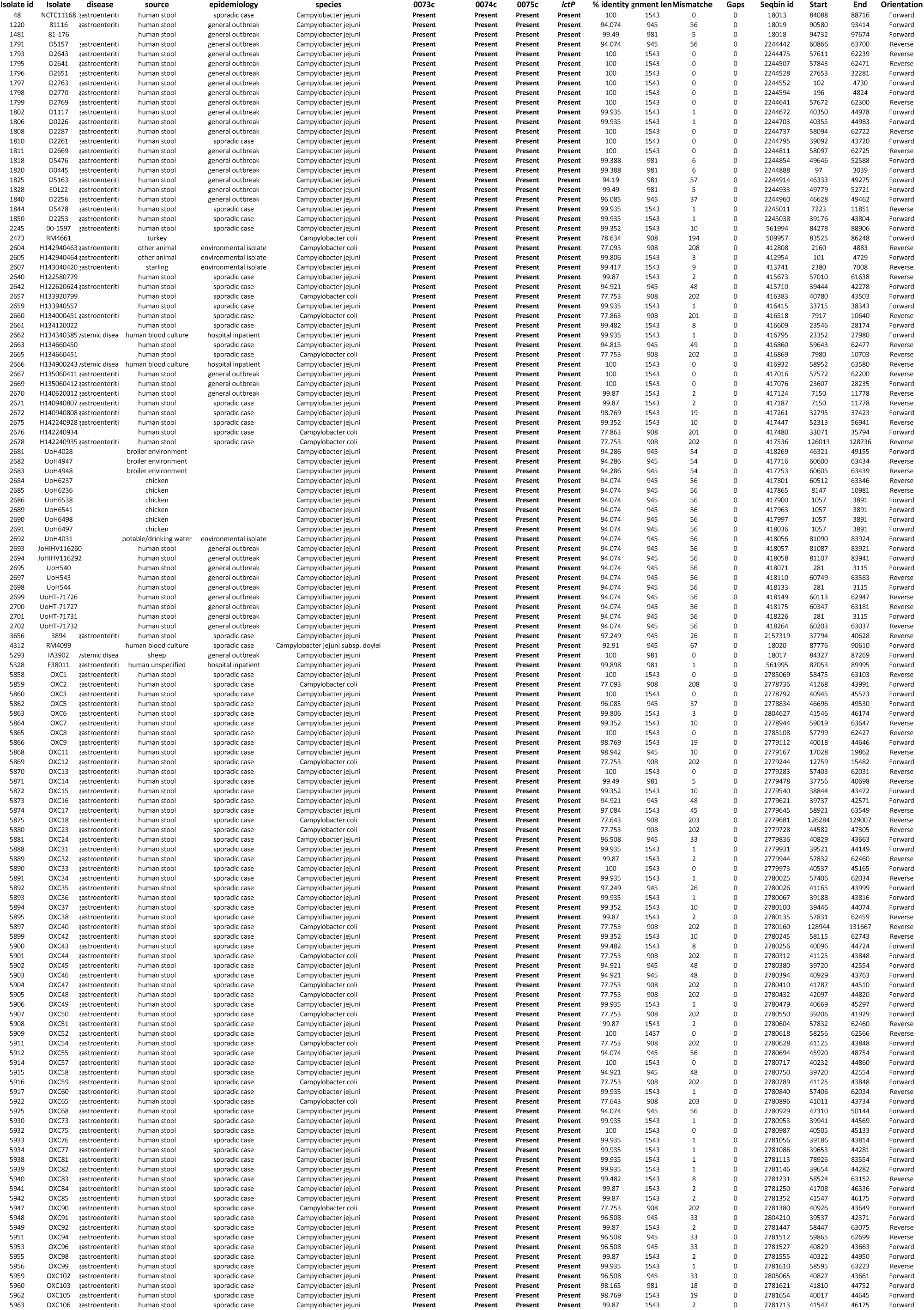
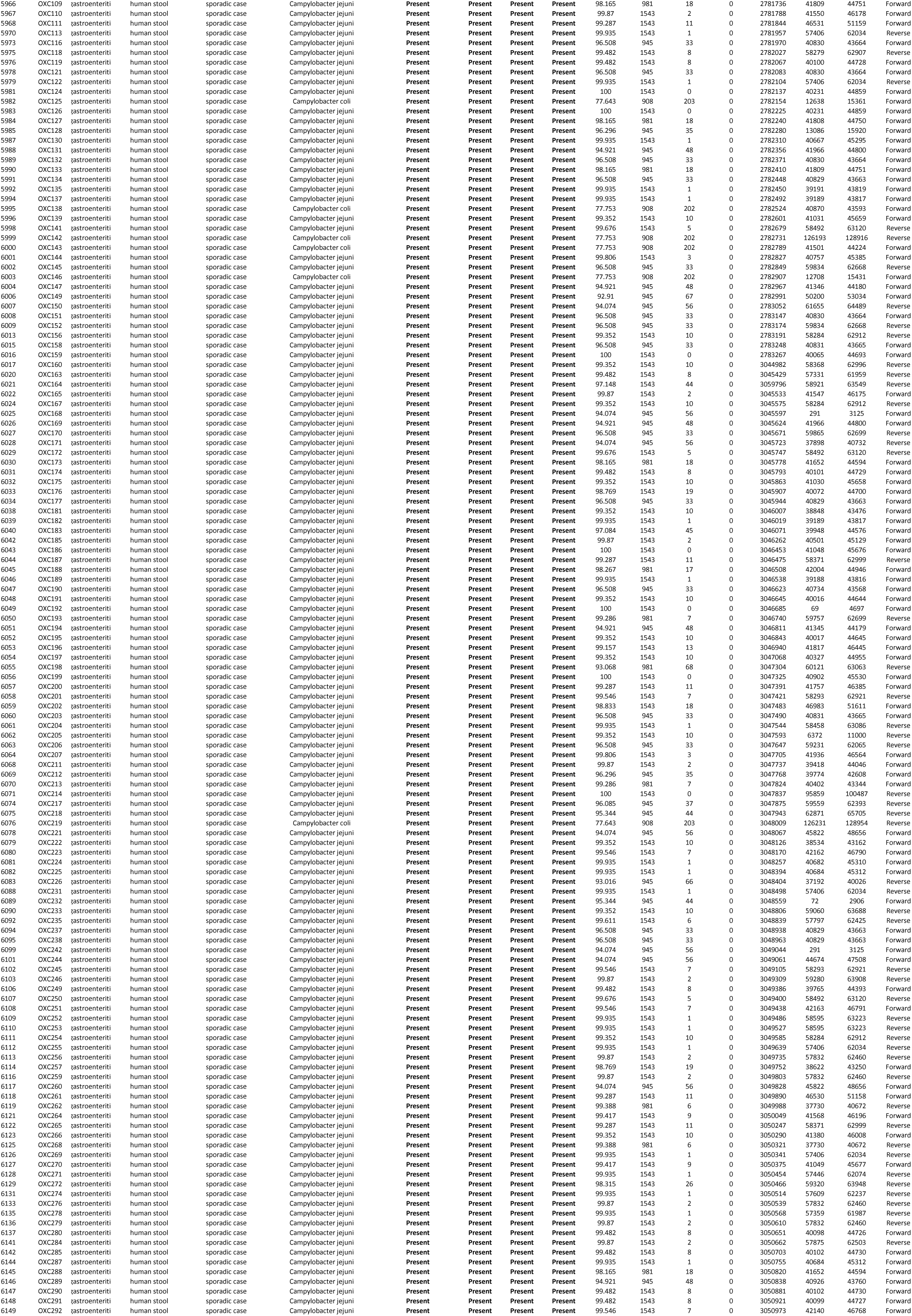
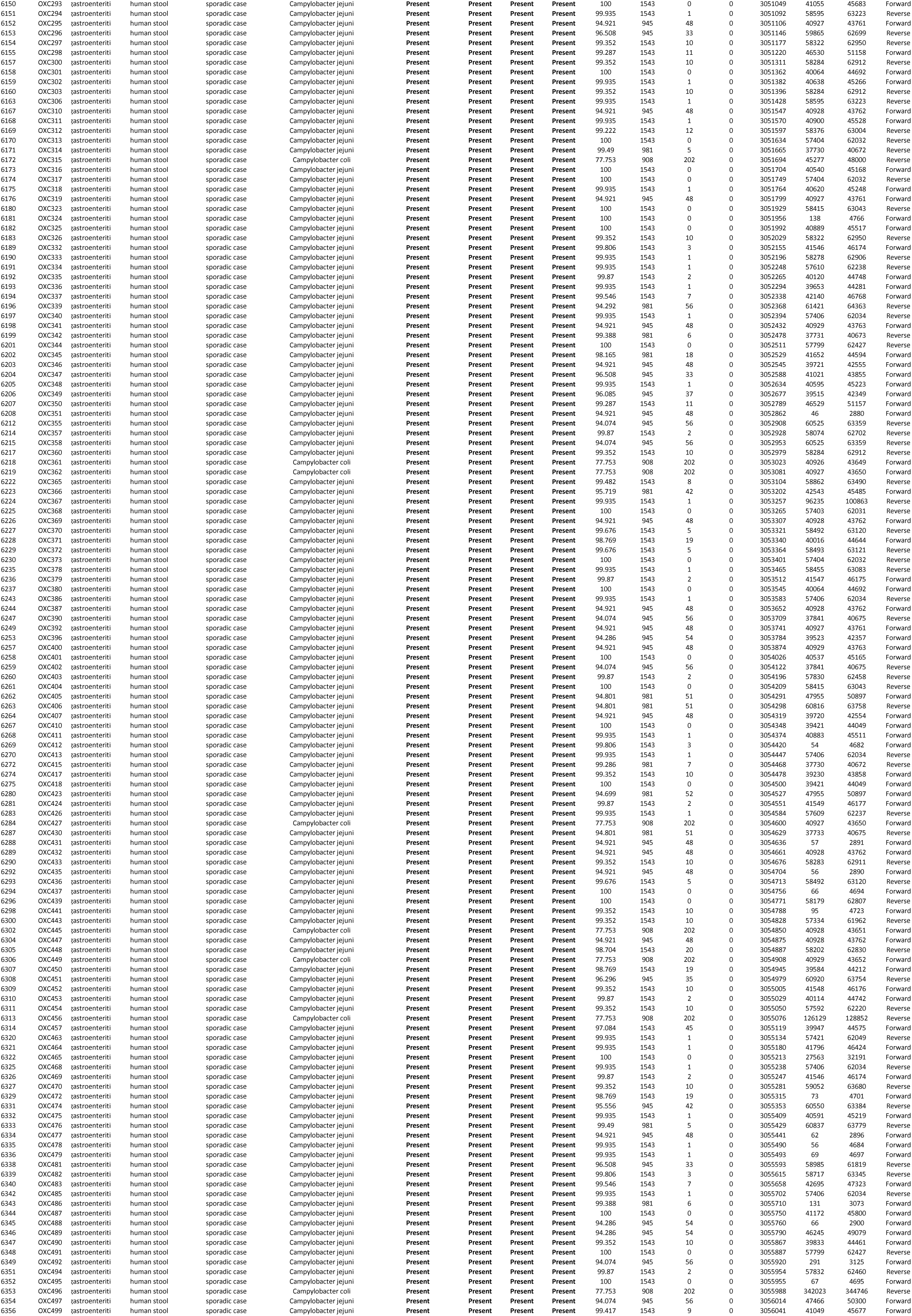
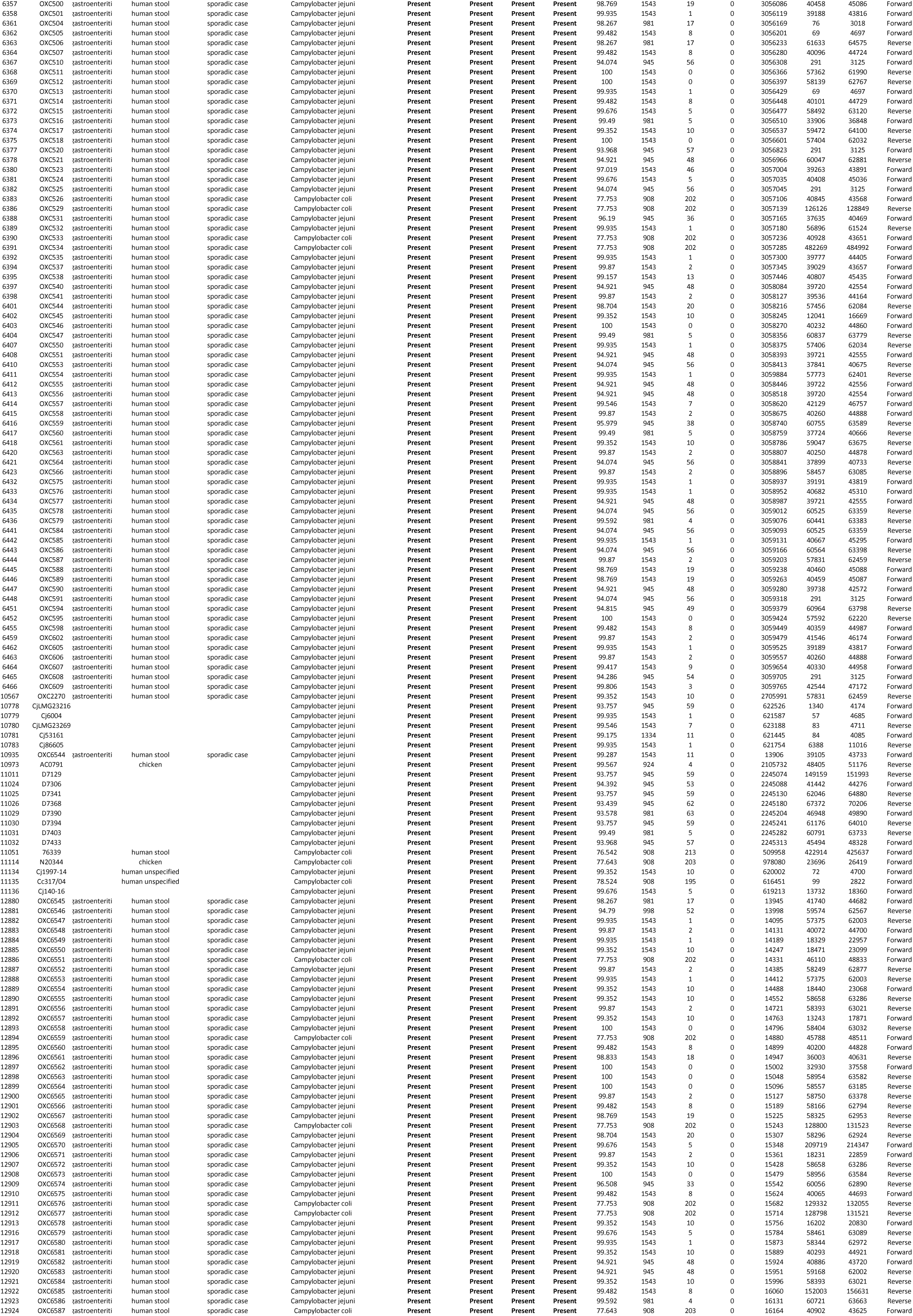
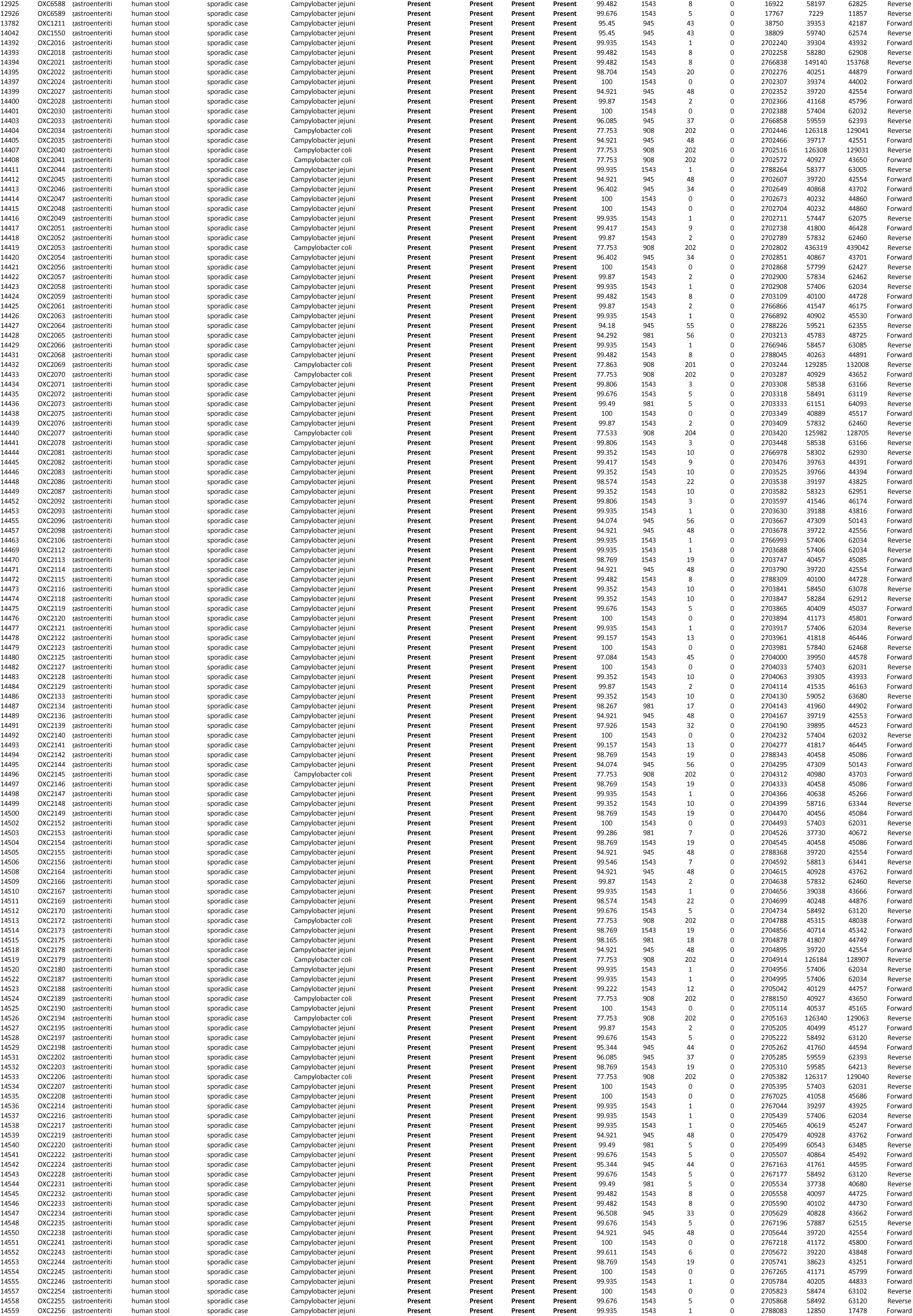
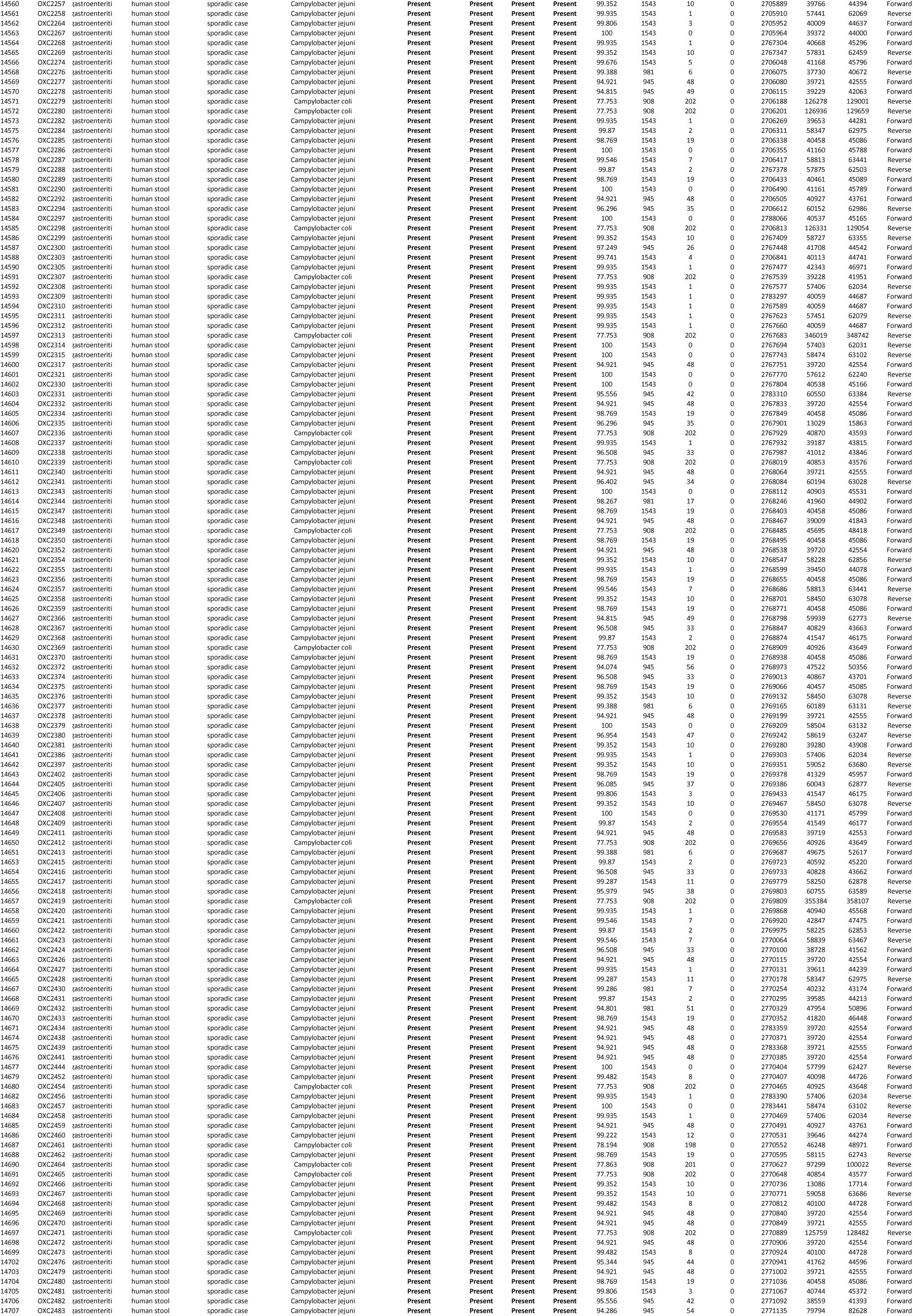
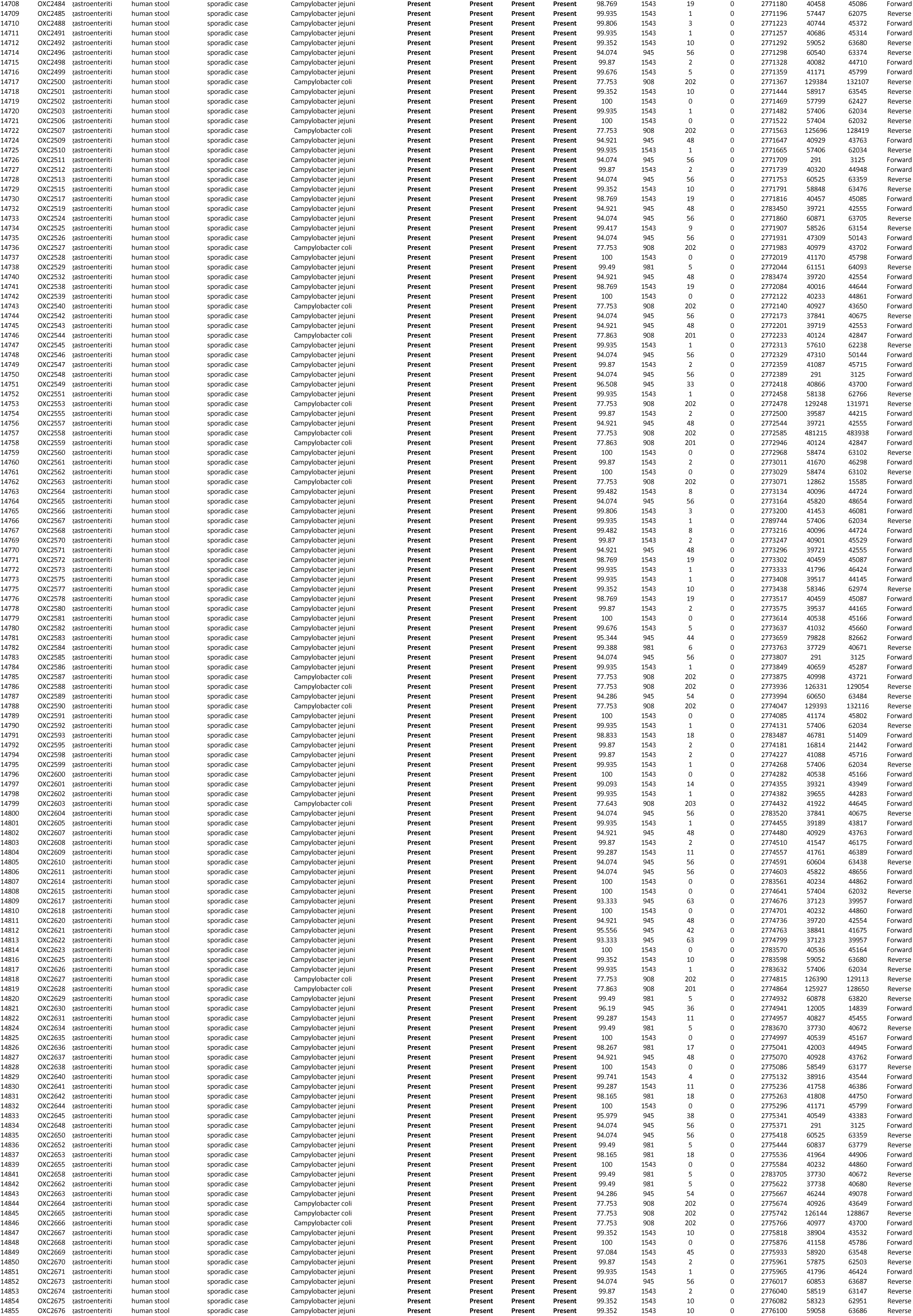
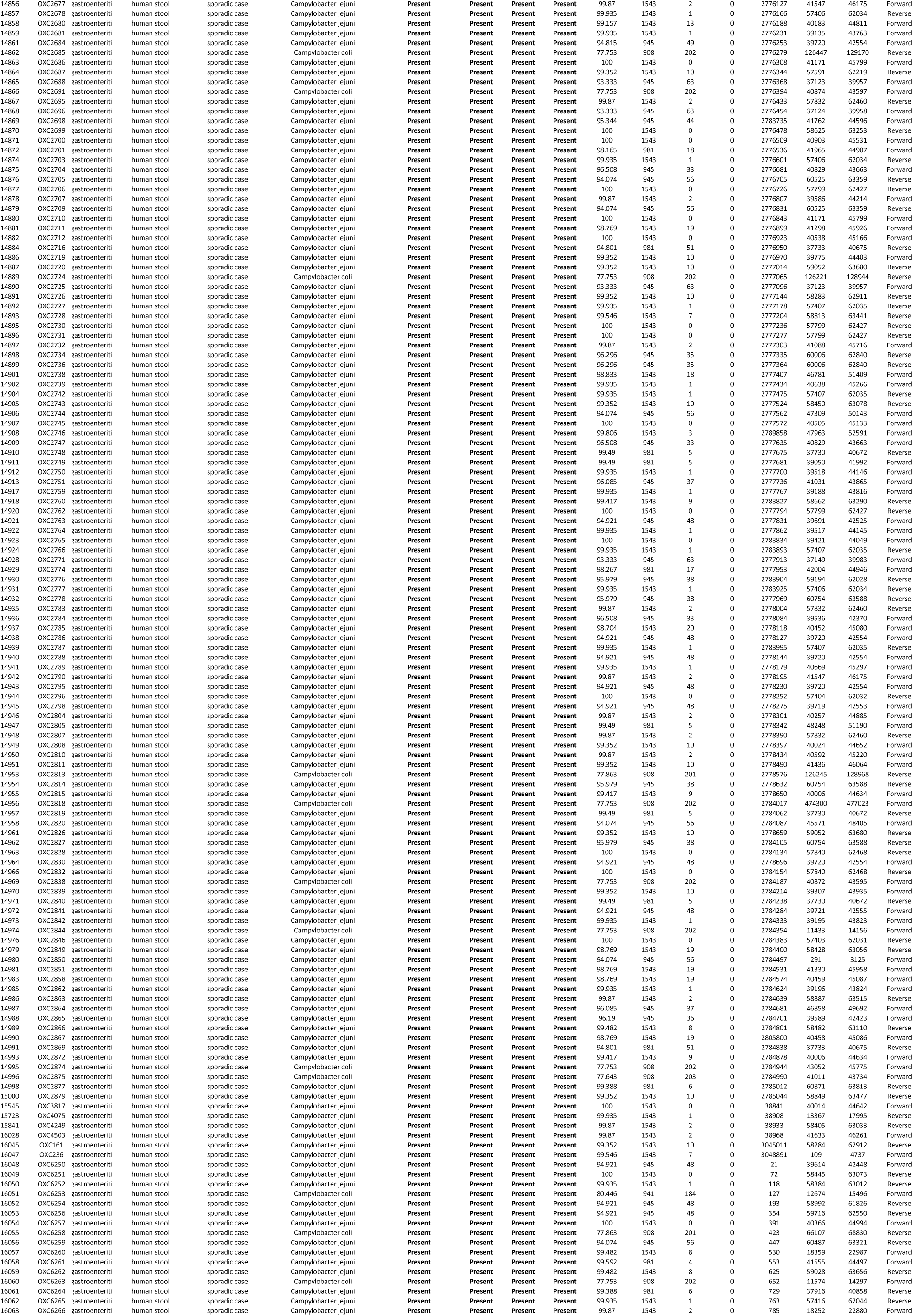
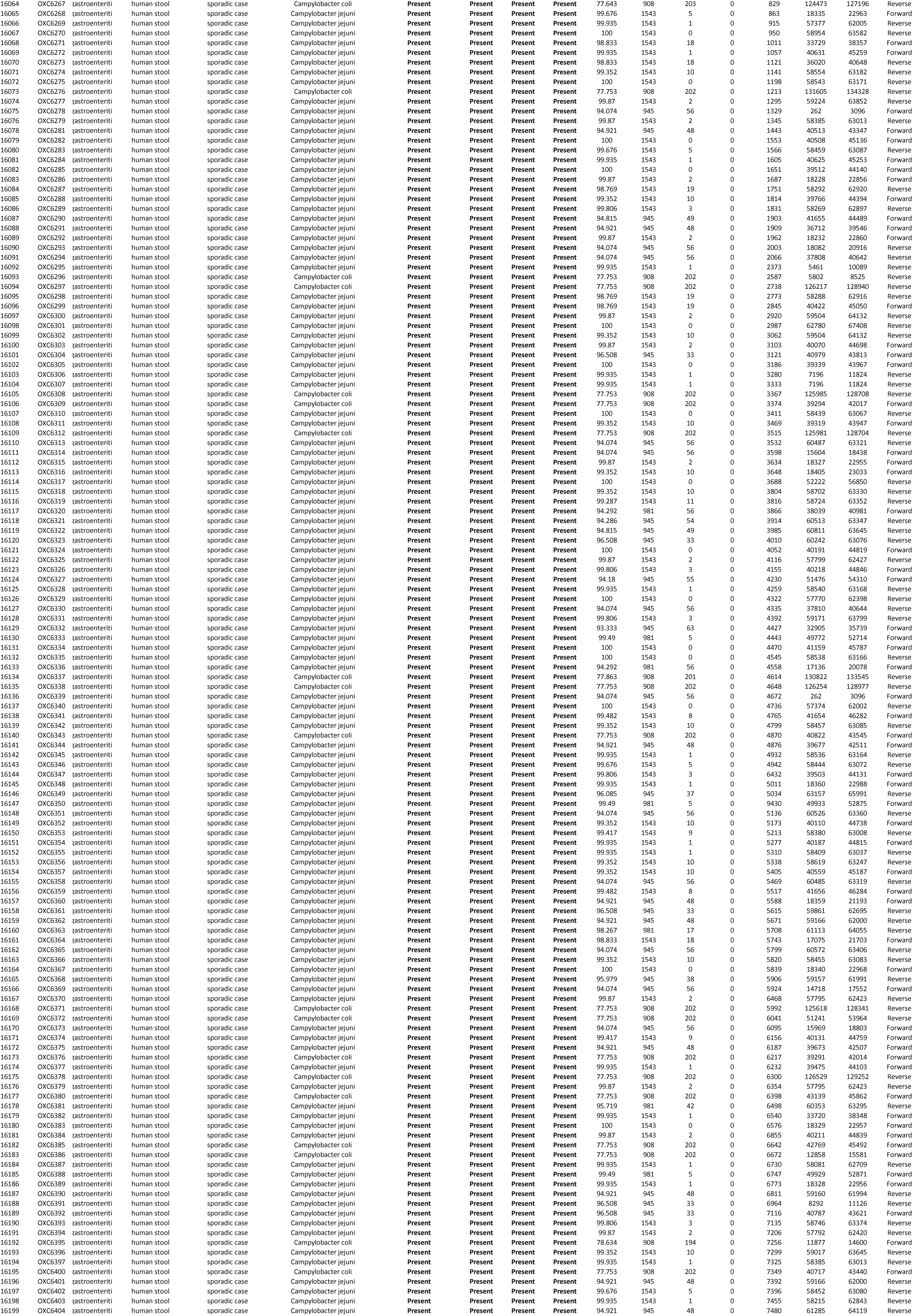
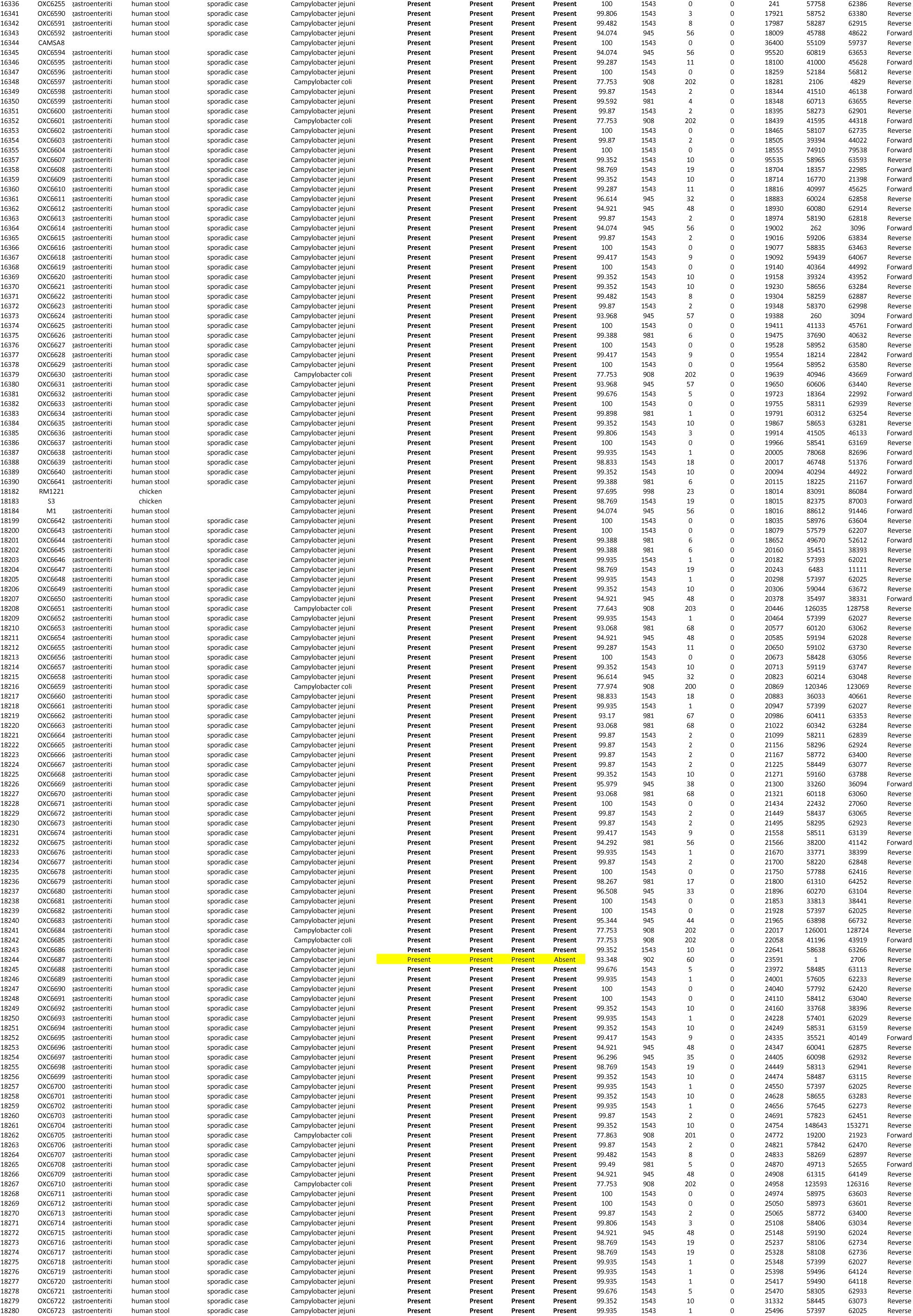
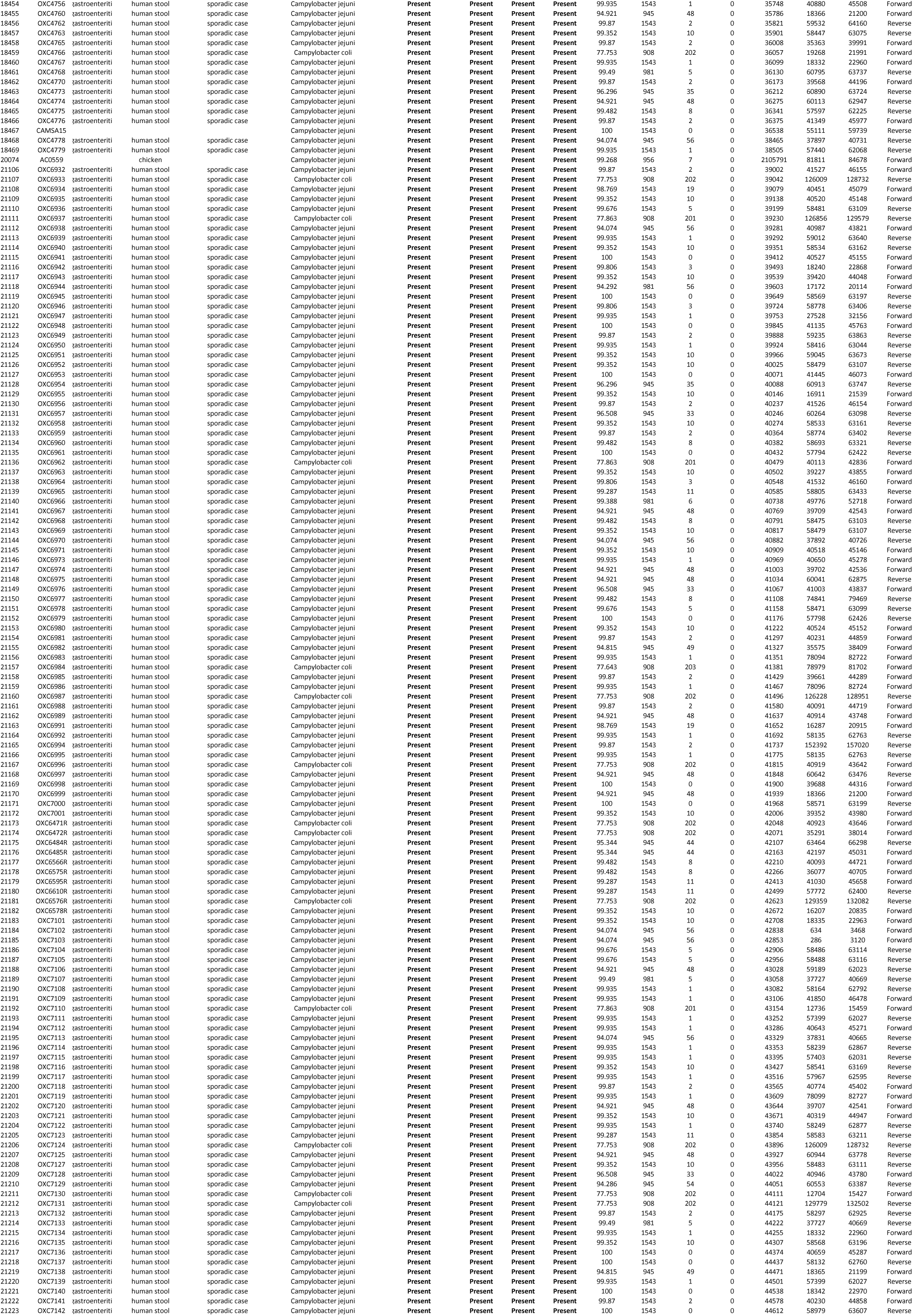
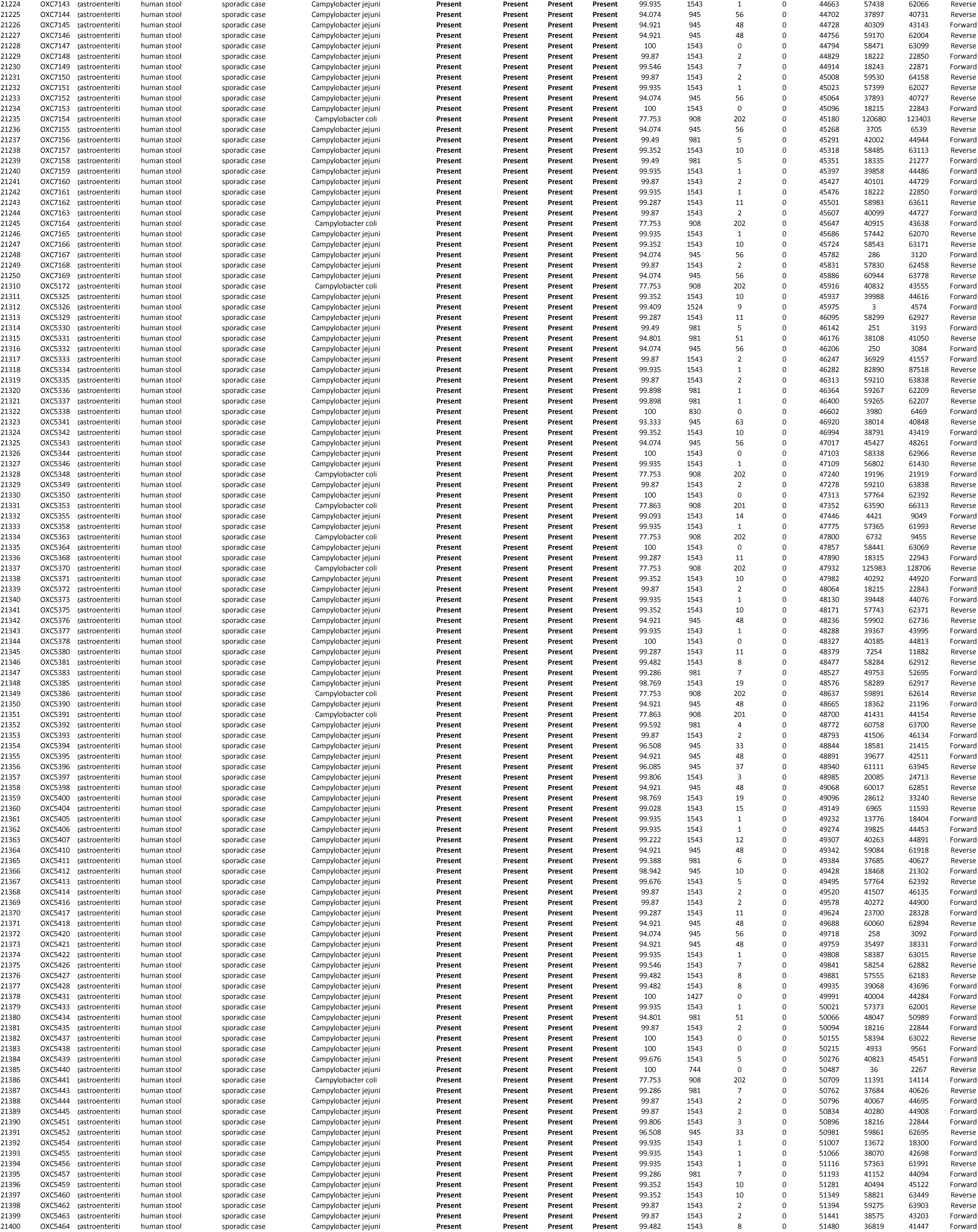
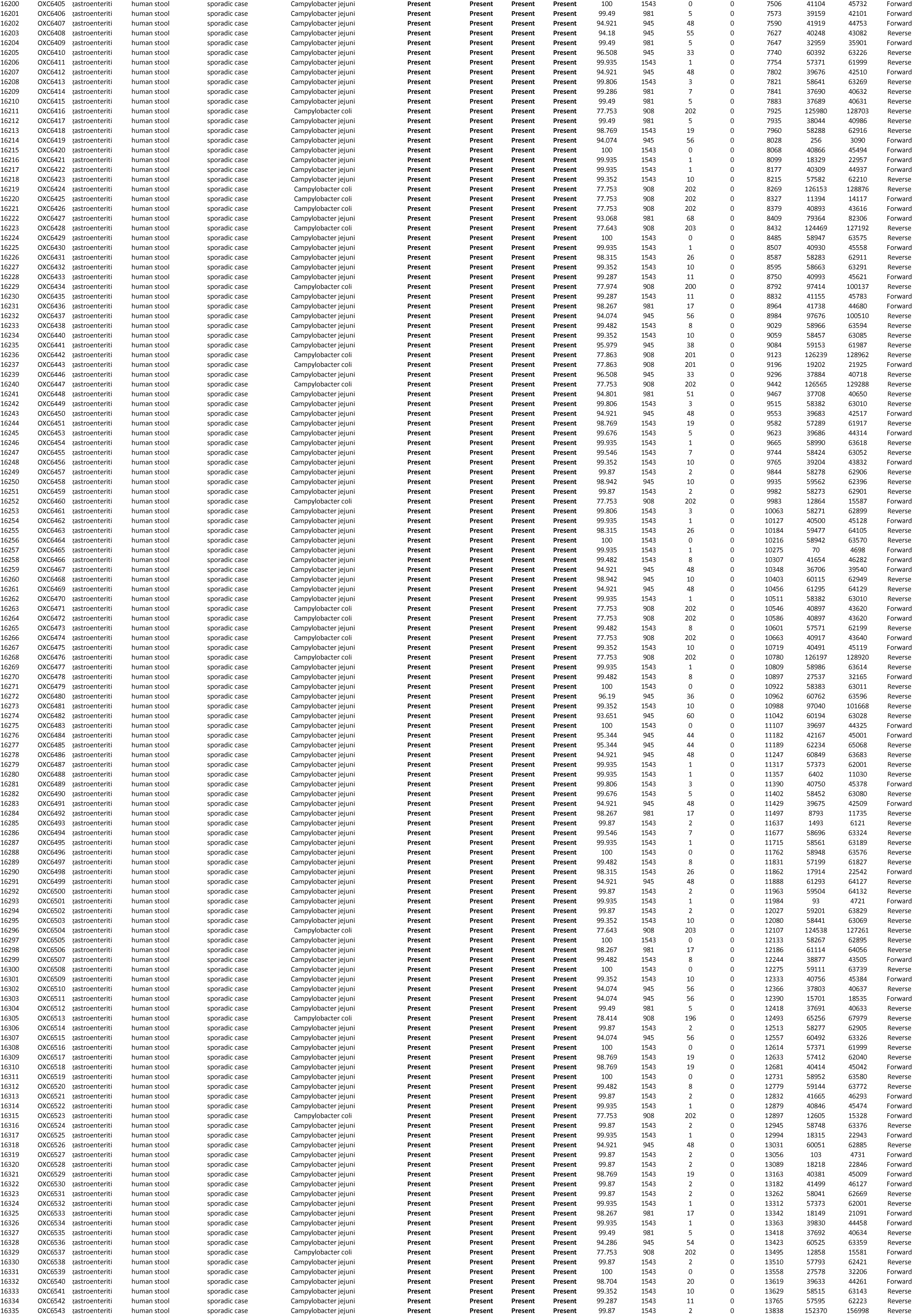
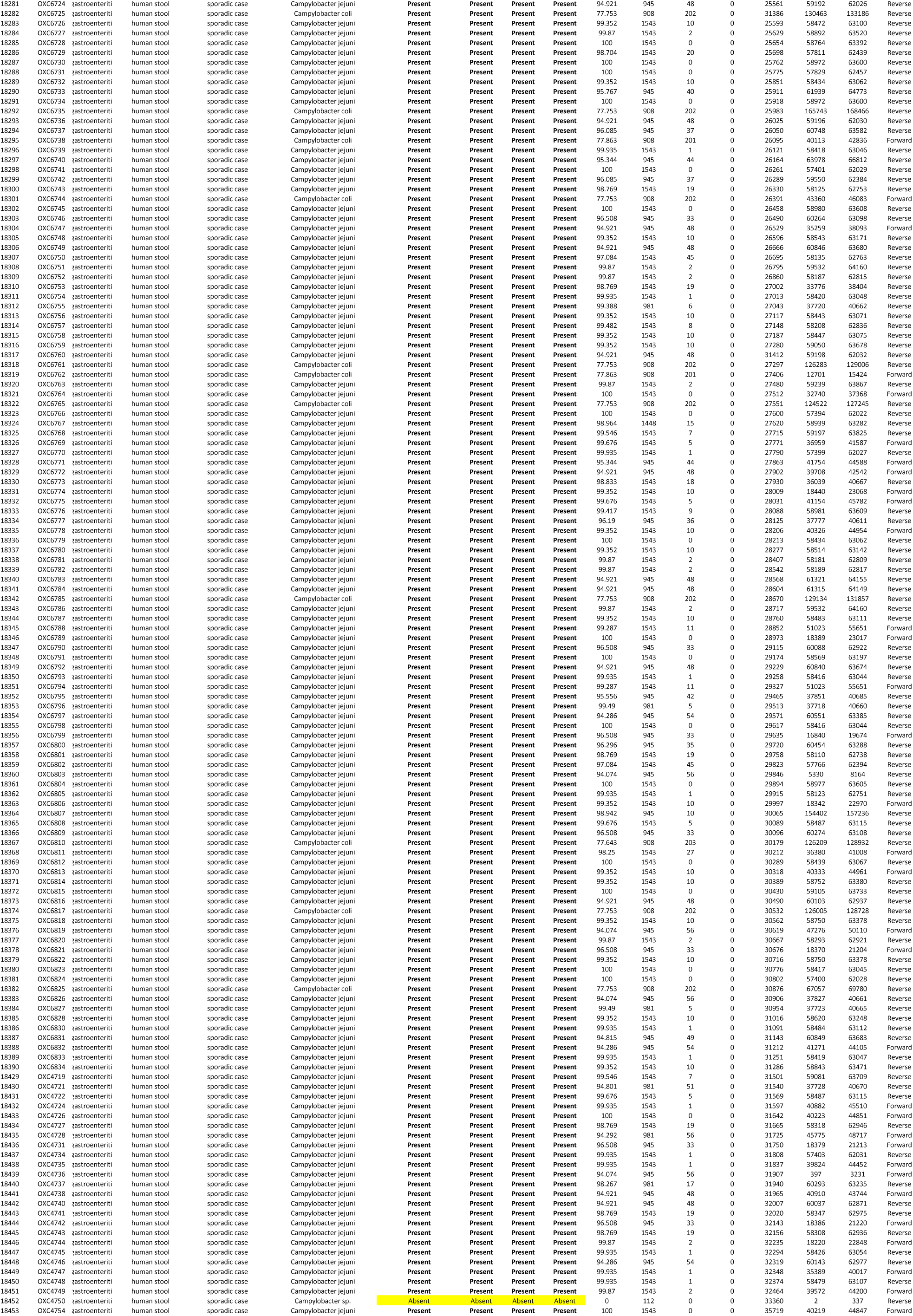

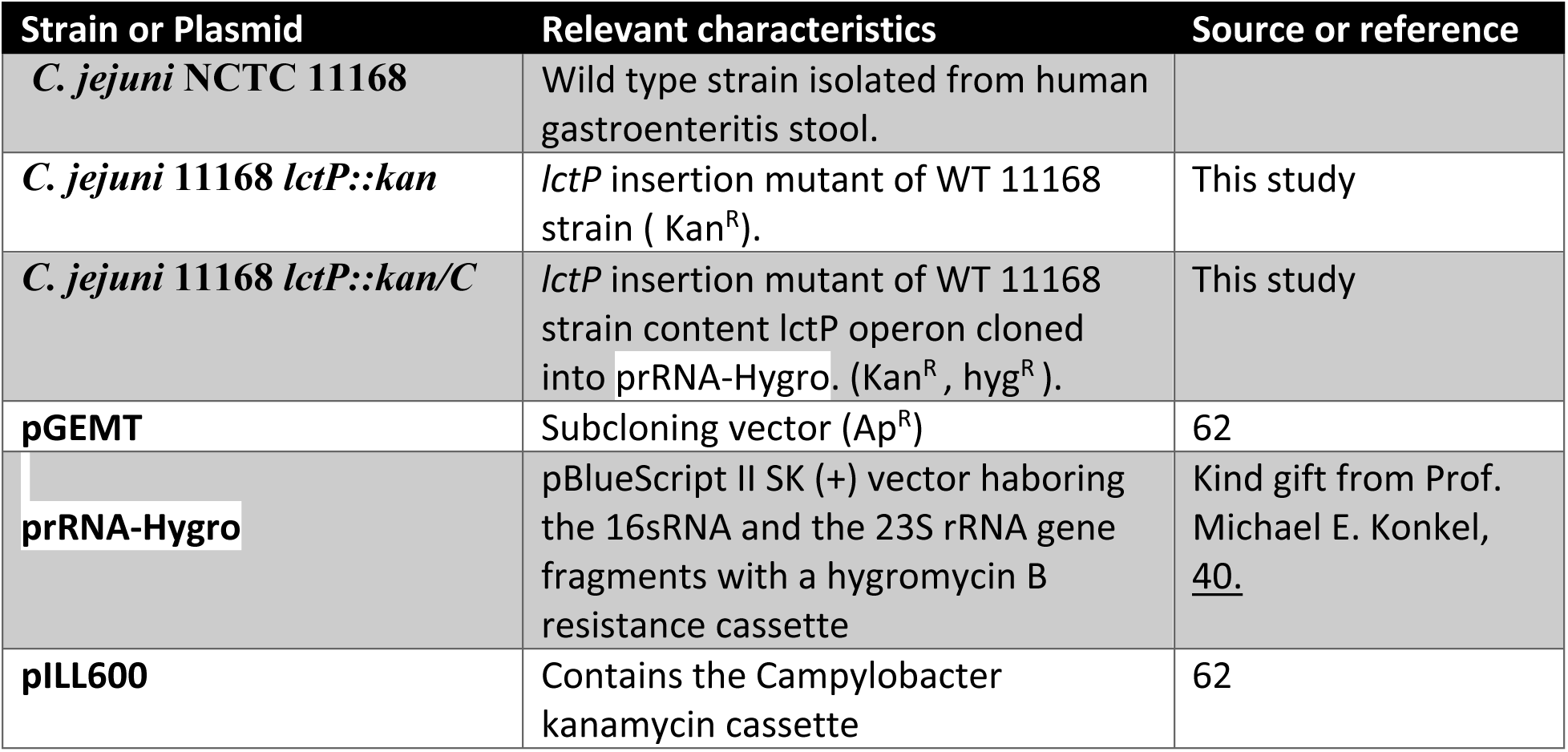
Table S2:

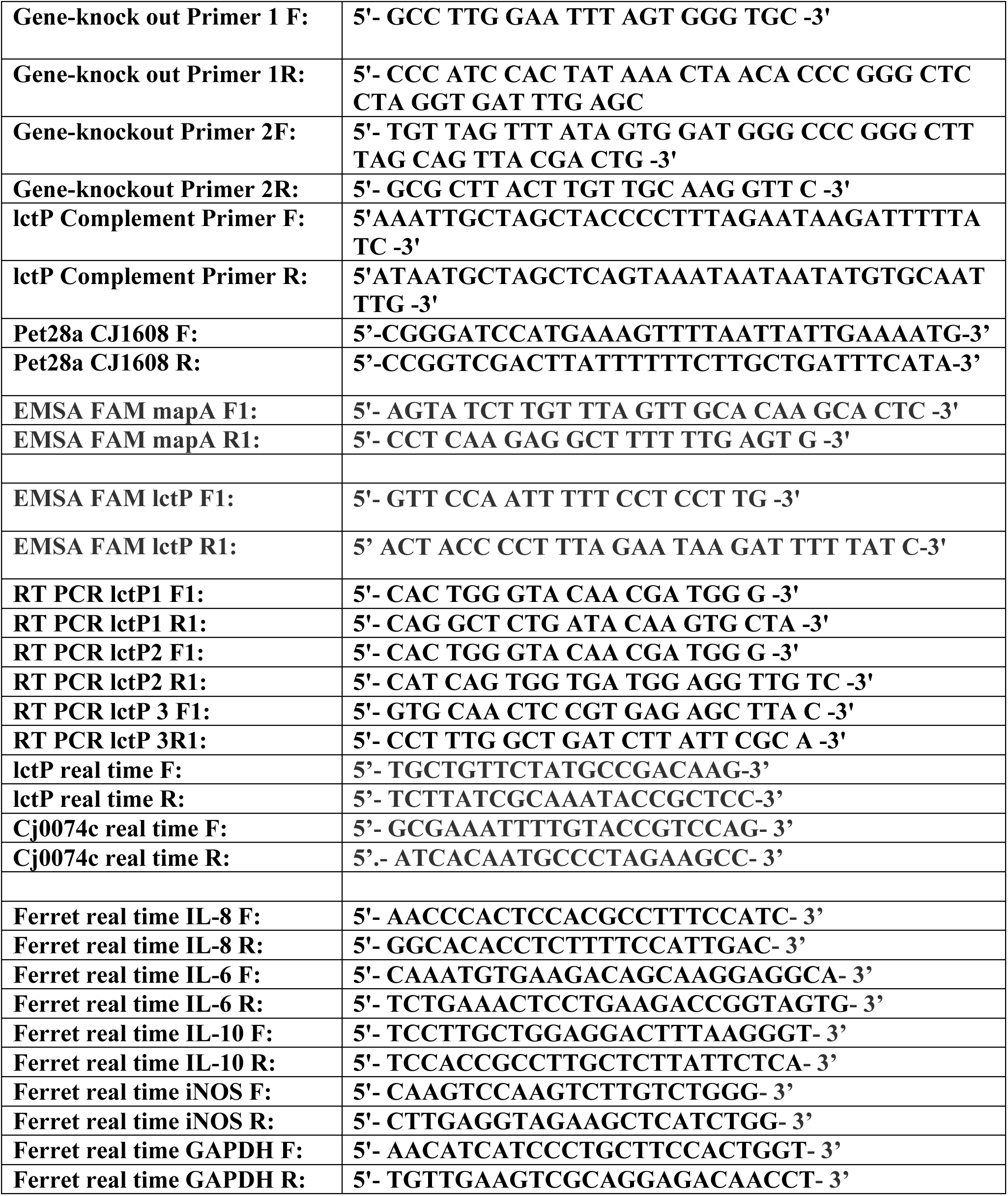
Table S3:

