## Supplemental Tables, Primers for "Gut metabolite L-lactate supports *Campylobacter jejuni* population expansion during acute infection"

**Table S2:**

| Strain or Plasmid | Relevant characteristics | Source or reference |
| --- | --- | --- |
| *C. jejuni* NCTC 11168 | Wild type strain isolated from human gastroenteritis stool. |  |
| *C. jejuni* 11168 *lctP::kan* | *lctP* insertion mutant of WT 11168 strain ( Kan^R^). | This study |
| *C. jejuni* 11168 *lctP::kan/C* | *lctP* insertion mutant of WT 11168 strain content lctP operon cloned into prRNA-Hygro. (Kan^R^ , hyg^R^ ). | This study |
| pGEMT | Subcloning vector (Ap^R^) | 62 |
| prRNA-Hygro | pBlueScript II SK (+) vector haboring the 16sRNA and the 23S rRNA gene fragments with a hygromycin B resistance cassette | Kind gift from Prof. [Michael E. Konkel](https://pubmed.ncbi.nlm.nih.gov/?term=Konkel%20ME%5BAuthor%5D), 40. |
| pILL600 | Contains the Campylobacter kanamycin cassette | 62 |

**Table S3:**

| **Gene-knock out Primer 1 F:** | **5'- GCC TTG GAA TTT AGT GGG TGC -3'** |
| --- | --- |
| **Gene-knock out Primer 1R:** | **5'- CCC ATC CAC TAT AAA CTA ACA CCC GGG CTC CTA GGT GAT TTG AGC** |
| **Gene-knockout Primer 2F:** | **5'- TGT TAG TTT ATA GTG GAT GGG CCC GGG CTT TAG CAG TTA CGA CTG -3'** |
| **Gene-knockout Primer 2R:** | **5'- GCG CTT ACT TGT TGC AAG GTT C -3'** |
| **lctP Complement Primer F:** | **5'AAATTGCTAGCTACCCCTTTAGAATAAGATTTTTATC -3'** |
| **lctP Complement Primer R:** | **5'ATAATGCTAGCTCAGTAAATAATAATATGTGCAATTTG -3'** |
| **Pet28a CJ1608 F:** | **5’-CGGGATCCATGAAAGTTTTAATTATTGAAAATG-3’** |
| **Pet28a CJ1608 R:** | **5’-CCGGTCGACTTATTTTTTCTTGCTGATTTCATA-3’** |
| **EMSA FAM mapA F1:** | **5'- AGTA TCT TGT TTA GTT GCA CAA GCA CTC -3'** |
| **EMSA FAM mapA R1:** | **5'- CCT CAA GAG GCT TTT TTG AGT G -3'** |
| **EMSA FAM lctP F1:** | **5'- GTT CCA ATT TTT CCT CCT TG -3'** |
| **EMSA FAM lctP R1:** | **5’ ACT ACC CCT TTA GAA TAA GAT TTT TAT C-3'** |
| **RT PCR lctP1 F1:** | **5'- CAC TGG GTA CAA CGA TGG G -3'** |
| **RT PCR lctP1 R1:** | **5'- CAG GCT CTG ATA CAA GTG CTA -3'** |
| **RT PCR lctP2 F1:** | **5'- CAC TGG GTA CAA CGA TGG G -3'** |
| **RT PCR lctP2 R1:** | **5'- CAT CAG TGG TGA TGG AGG TTG TC -3'** |
| **RT PCR lctP 3 F1:** | **5'- GTG CAA CTC CGT GAG AGC TTA C -3'** |
| **RT PCR lctP 3R1:** | **5'- CCT TTG GCT GAT CTT ATT CGC A -3'** |
| **lctP real time F:** | **5’- TGCTGTTCTATGCCGACAAG-3’** |
| **lctP real time R:** | **5’- TCTTATCGCAAATACCGCTCC-3’** |
| **Cj0074c real time F:** | **5’- GCGAAATTTTGTACCGTCCAG- 3’** |
| **Cj0074c real time R:** | **5’.- ATCACAATGCCCTAGAAGCC- 3’** |
| **Ferret real time IL-8 F:** | **5'- AACCCACTCCACGCCTTTCCATC- 3’** |
| **Ferret real time IL-8 R:** | **5'- GGCACACCTCTTTTCCATTGAC- 3’** |
| **Ferret real time IL-6 F:** | **5'- CAAATGTGAAGACAGCAAGGAGGCA- 3’** |
| **Ferret real time IL-6 R:** | **5'- TCTGAAACTCCTGAAGACCGGTAGTG- 3’** |
| **Ferret real time IL-10 F:** | **5'- TCCTTGCTGGAGGACTTTAAGGGT- 3’** |
| **Ferret real time IL-10 R:** | **5'- TCCACCGCCTTGCTCTTATTCTCA- 3’** |
| **Ferret real time iNOS F:** | **5'- CAAGTCCAAGTCTTGTCTGGG- 3’** |
| **Ferret real time iNOS R:** | **5'- CTTGAGGTAGAAGCTCATCTGG- 3’** |
| **Ferret real time GAPDH F:** | **5'- AACATCATCCCTGCTTCCACTGGT- 3’** |
| **Ferret real time GAPDH R:** | **5'- TGTTGAAGTCGCAGGAGACAACCT- 3’** |
